# Compensating for a sensorimotor delay requires a predictor that convolves over a memory buffer of efference copies

**DOI:** 10.1101/2024.11.18.624125

**Authors:** Eric Maris

## Abstract

Effective motor control requires sensory feedback but is seriously complicated by the sensorimotor delay (SMD), which is the time delay between the state of the body at feedback generation and the arrival in the body’s muscles of the feedback-informed motor command. I describe and evaluate three SMD compensation mechanisms: gain scaling, the convolution predictor, and the Smith predictor. These mechanisms are implemented using control theory results for linear dynamical systems, which are well motivated for balance control. These mechanisms are investigated theoretically and by simulations of balance control, both free standing and while riding a bicycle. I demonstrate that compensating for a SMD requires a convolution predictor, which involves a convolution over a memory buffer of efference copies and an initial condition obtained from a state observer that is based on a delayed-input forward model. The performance of a convolution predictor does not crucially depend on its exact computational implementation because a similar performance is obtained with an approximate convolution using a boxcar kernel. I also demonstrate that gain scaling is an effective SMD compensation mechanism but is not sufficient to compensate for a neurobiological SMD. Finally, I demonstrate that the Smith predictor is an ineffective and neurobiologically implausible SMD compensation mechanism for an unstable mechanical system.

## Introduction

The central nervous system (CNS) controls the body via motor commands, and this usually requires sensory feedback about the state of the body (i.e., its position and velocity). Adjusting motor commands based on sensory feedback is complicated by the delay between the sensory feedback (informing us about the state of our body) and the effect of the feedback-based motor commands on the body. This is called the sensorimotor delay (SMD). For example, in balance control, the motor commands that keep the body upright lag the sensory feedback about the body’s state relative to gravity. Motor control deteriorates as the SMD is artificially increased [1-3], but repeated exposure to such imposed delays (up to 430 ms.) results in improvements in motor control [4-6]. This suggests that the CNS implements computational mechanisms that compensate for the SMD.

In this paper, I describe and evaluate three SMD compensation mechanisms. I evaluate these mechanisms both theoretically and by simulating their performance in balancing two unstable mechanical systems, a freely standing person and a rider-bicycle combination. The SMD compensation mechanisms are implemented using control theory results for linear dynamical systems [7, 8]. Although mechanical systems involving the human body have nonlinear equations of motion (EoM), for balance control, a linear approximation of these nonlinear EoM is accurate. This is because a balancing body (standing person, rider, bicycle) most of the time stays close to the upright position, which is an unstable fixed point of the nonlinear mechanical system. The linear approximation at this unstable fixed will be accurate as long as the body stays close to it.

Control theory allows to formulate SMD compensation mechanisms in terms of a CNS-internal dynamical system (the computational system) that controls the mechanical system (the plant). This mechanical system is the human body plus the objects attached to it (e.g., a bicycle), and its dynamics depends on the forces that act on it, including gravity and the centripetal force. Control is specified in terms of an optimality criterion/objective. In the time domain, this criterion is usually expressed in terms of a loss function, and for balance control this loss function depends on the difference between the body’s orientation and the earth-vertical axis. In the Laplace domain, the optimality criterion is expressed in terms of the poles of transfer functions. The best-known type of control is linear-quadratic-Gaussian (LQG) control [9, 10], which contains all the ingredients that are needed as a formal background for this paper. Crucially, although I have used LQG in my simulations, the main concepts in this paper do not depend on the exact mathematical specification of the optimality criterion.

In the absence of a SMD, LQG-optimal control for a linear mechanical system is realized by a control action that obeys the separation principle. Under this principle, the optimal control action *u* is of the following linear type: 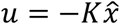 , in which −*K* is an output gain and 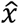 is an estimate of the mechanical system’s state. The separation principle involves that the optimal control action *u* is obtained by two separate optimizations: (1) find the optimal (most accurate) state estimate 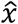 for a fixed output gain −*K*, and (2) find the optimal output gain −*K* for a state estimate 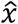. Under the LQG assumptions, the optimal state estimate 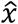 is the output of a Kalman filter that takes as its input the sensory feedback *y* and the control action *u*, and the optimal output gain −*K* is the Linear Quadratic Regulator (LQR).

I assume that, also in the presence of a nonzero SMD, the CNS obeys the separation principle: one CNS component is responsible for computing the optimal state estimate 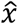 , and another component for computing the optimal control action 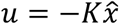 Corresponding to these two components, there are also two types of SMD compensation mechanisms: (1) state prediction, and (2) output gain scaling. State prediction involves that the state estimate 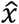 is replaced by a state prediction. Two types of state prediction have been proposed: (1) finite spectrum assignment (FSA) [11-17], and (2) the Smith predictor [11, 18-20]. Output gain scaling [21-23] involves downscaling of the output gain, and this is motivated quantitatively in the Laplace domain: by downscaling the output gain, the phase margin of the implied open-loop system is evaluated at a lower frequency, which has a smaller SMD-induced phase lag and therefore a larger phase margin.

All three mechanisms have their origin in engineering, but they generate useful ideas about the computations that are likely to be performed by the CNS. A very important idea is that the CNS uses internal models for motor control [24, 25]. Different internal models have been proposed, and here I focus on the forward and the sensory model: the forward model captures the dynamics of the mechanical system (including its dependence on the muscular input forces), and the sensory model captures the information in the sensory feedback about the mechanical system’s state. Also an inverse model has been proposed [26-28], but its function is identical to that of the output gain −*K*, and I will therefore not use this term.

FSA and the Smith predictor differ with respect to the force input to their forward model: in the FSA forward model, the input force is delayed, whereas in the Smith predictor’s forward model, it is non-delayed. These different forward models have implications for the CNS’ computations: (1) FSA is a state predictor that depends on a memory buffer of efference copies, and (2) the Smith predictor is a forward state estimator with a correction term that depends on a memory buffer of state estimates.

Finite spectrum assignment (FSA) is a poor label for a technique that serves as an inspiration for CNS computations. The label FSA refers to the Laplace domain formulation of the technique, whereas the time domain formulation can be related much easier to CNS computations. In its time domain formulation, FSA is a state predictor produced by convolution. This differs from the Smith predictor, which is a state estimator combined with a term that is intended to correct for the SMD; it is the correction term that justifies the label predictor. In the following, instead of FSA, I will use the label convolution predictor.

The control theory literature on delay compensation is dominated by theoretical analyses and examples with sensory feedback that is a perfect copy of the state variables. Such feedback is called full-state and noise-free. Full-state noise-free feedback is unlikely to be observed in a neurobiological system, and therefore it is important to investigate whether SMD compensation mechanisms can control an unstable mechanical system using sensory feedback in which the states are only partially observable (denoted as incomplete feedback) and noisy. In this paper, I model muscle spindle proprioceptive feedback based on results from sensory neurophysiology: muscle spindle primary afferents scale with exafferent acceleration feedback [29-31], which is incomplete [10]. LQG-optimal control of a mechanical system using incomplete and noisy feedback requires an optimal state estimator (observer). This optimal observer is the Kalman filter [32, 33], which is a CNS-internal dynamical system that can be implemented as a recurrent neural network [10].

I will demonstrate that the convolution predictor is the only plausible SMD compensation mechanism. Although standing balance control does not require a compensation mechanism under a neurobiologically plausible SMD, bicycle balance control is only possible using the convolution predictor. The Smith predictor is an ineffective mechanism for an unstable mechanical system, and gain scaling is not sufficiently effective for bicycle balance control. The convolution predictor requires a memory buffer of efference copies plus a state observer that is based on a delayed-input forward model. Effective SMD compensation using the convolution predictor does not require the computationally hard convolution; balance control performance is almost identical if a grossly simplified approximate convolution is used. None of the results are specific for the incomplete exafferent acceleration feedback because qualitatively similar results were obtained using full-state feedback.

## Results

### A model for a standing person’s body: the double compound inverted pendulum

For a concrete description of the SMD compensation mechanisms, it is useful to start from a mechanical system that models the musculoskeletal properties of a standing person’s body. For that purpose, I use the double compound inverted pendulum (DCIP), which is depicted in **Error! Reference source not found**.A. The DCIP has two parts, of which one represents the lower and the other the upper body. The DCIP has two joints, one at the ankle and one at the hip. The angular position of the lower and the upper body relative to gravity are denoted by, respectively, *θ*_1_ and *θ*_2_. The muscular forcing torque at the ankle and the hip joint are denoted by, respectively, *z*_1_ and *z*_2_. I will denote these angular positions and forcing torques as vectors ***θ*** = [*θ*_1_, *θ*_2_]^*t*^ = [*θ*_1_; *θ*_2_] and ***z*** = [*z*_1_, *z*_2_]^*t*^ = [*z*_1_; *z*_2_]. These variables are all functions of time, but I will suppress their argument *t* whenever the context allows.

The movements of the DCIP are fully specified by its EoM, which express the angular acceleration 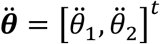 as a function of angular position ***θ***, angular velocity 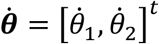 , and forcing torque ***z***. EoM of multibody mechanical systems are usually derived using the method of Euler-Lagrange, which is straightforward but tedious. Here, I only give the result of this derivation, and I suppress a lot of detail:

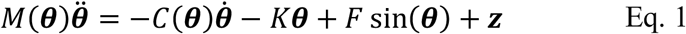

The active dynamics of the DCIP depend on the input/forcing torque ***z***, and the passive dynamics depend on the matrices *M*(***θ***) (mass moment of inertia), *C*(***θ***) (damping matrix), *K* (stiffness matrix), and *F* (matrix specifying the gravitational torque). Eq. 1 is nonlinear because of sin(***θ***), and because of the way the matrices *M*(***θ***) and *C*(***θ***) depend on ***θ*** (via sines and cosines).

The matrices *M*(***θ***), *C*(***θ***), *K*, and *F* depend on parameters that determine, respectively, the system’s mass moment of inertia, the joints’ damping and stiffness, and the gravitational torque. The values of these parameters are crucial for the dynamic properties of the DCIP. For instance, the DCIP is only unstable if the stiffness of the joints is less than a gravity-dependent constant (the critical stiffness) that determines the gravitational torque per unit of lean angle. Fortunately, the values of the crucial parameters have been measured. For instance, several studies have demonstrated that the ankle joint stiffness is less than the critical stiffness, and that this is due to the compliance of the Achilles tendon [34-37].

To express 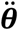 as a function of ***θ***, 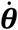 and ***z***, the left and right side of Eq. 1 must be premultiplied by *M*(***θ***)^−1^. Usually, ***θ*** and 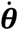 are combined in the vector 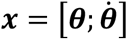 , which is called the state of the mechanical system. The EoM can then be written as a first order nonlinear differential equation for the state ***x***, which I will denote as 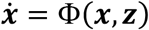 note that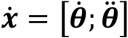. This differential equation is usually called the EoM in state space form and is described in more detail in the Methods section.

In the neighborhood of ***x*** = **0**, the nonlinear differential equation 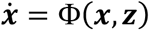 can be approximated by a linear differential equation of which the coefficient matrices are obtained as the Jacobian of Φ(***x***, ***z***) and evaluated at ***x*** = **0** and ***z*** = **0**. Note that 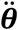 depends linearly on ***z***. This is also called a Taylor approximation of Φ(***x***, ***z***). This linear approximation is accurate for a standing person that is almost stationary (i.e.,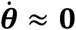) and stays close to the upright position (i.e., ***θ*** ≈ **0**). It can be written as follows:

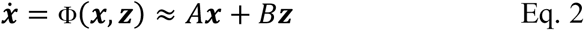

in which *A* and *B* are the Jacobian matrices of Φ(***x***, ***z***) with respect to, respectively, ***x*** and ***z***, and evaluated at ***x*** = **0** and ***z*** = **0**.

### Overview of the SMD compensation mechanisms

The three SMD compensation mechanisms are depicted in Fig. 1B-D. Fig. 1B shows a block diagram of closed-loop feedback control with SMD compensation by means of gain scaling. The diagram involves a mechanical (in red), a sensory (in green), a computational (blue, black and purple elements; see below), and a motor output system (in black) whose input is modulated by gain scaling (in purple). The computational system models the CNS as a neural network that (1) receives the sensory feedback ***y*** in its input level (green rectangle), (2) has an intermediate level with recurrent connections (blue rectangle), and (3) produces output ***u*** via two feedforward levels (purple and black rectangle). The black-and-white box depicts the delay between output ***u*** and the mechanical system state ***x*** (motor delay, MD), and the green-and-white box depicts the delay between ***x*** and the sensory feedback ***y*** (sensory delay, SD). The SMD is equal to MD plus SD.

**Fig. 1:**
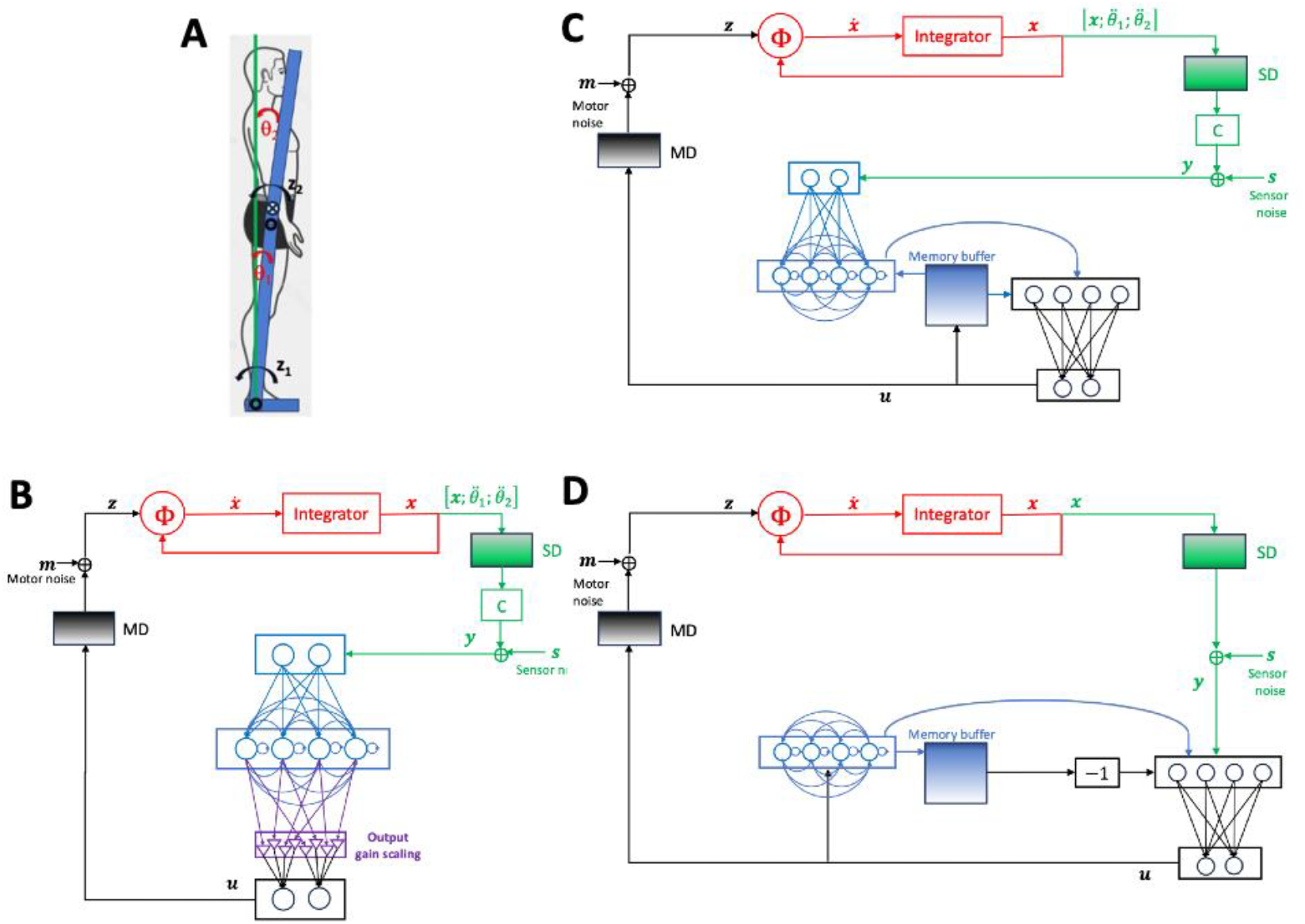
Three SMD compensation mechanisms applied to standing balance control. (A) A standing body with a superimposed double compound inverted pendulum rotating about the ankle and the hip, used to model standing balance control. See text for an explanation of the symbols. (B) Block diagram of closed-loop feedback control with SMD compensation by means of gain scaling (see text). (C) Block diagram of closed-loop feedback control with SMD compensation by means of the convolution predictor (see text). (D) Block diagram of closed-loop feedback control with SMD compensation by means of the Smith predictor (see text).

Fig. 1C shows a block diagram of closed-loop feedback control with SMD compensation by means of the convolution predictor. The main difference with the gain scaling in Fig. 1B is the role of the memory buffer of efference copies, which is depicted as a blue-and-white box. This memory buffer is used to store input for the computational and the motor output system.

Fig. 1D shows a block diagram of closed-loop feedback control with SMD compensation by means of the Smith predictor. The blue-and-white box now depicts a memory buffer of state estimates (produced by the recurrently connected level in blue) that stores input for the motor output system.

In Fig. 1B and 1C, the sensory system (in green) maps the vector 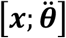 onto sensory variables, adds noise ***s*** and feeds the resulting feedback ***y*** into the computational system. By including the angular acceleration 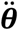 in the sensory feedback, I deviate from most of the existing literature on closed-loop feedback control, in which the sensory feedback only pertains to the state variable ***x***. This is because I use a sensory model for muscle spindle proprioceptive feedback, and sensory neurophysiology studies are well in line with the assumption that primary spindle afferents predominantly encode acceleration [38-41]. Angular acceleration sensory feedback can be modeled as incomplete state feedback and therefore requires a state observer [10].

The role of the sensory feedback is different for the Smith predictor and the two other mechanisms: the Smith predictor uses the sensory feedback to correct the state estimate such that it turns into a state prediction, whereas the other mechanisms use the sensory feedback to update the state estimate. To correct the state estimate in the Smith predictor, the sensory feedback must be full-state and noise-free (i.e., ***y***(*t*) = ***x***(*t* − *SD*)) and therefore Fig. 1D does not contain the matrix *C* that maps the state variables on the sensory feedback (or, equivalently, *C* is the identity matrix).

### A computational system for the scenario without SMD

My interest is in how the computational system can control the mechanical system in the presence of a SMD. It is useful to first describe this computational system for the scenario without SMD. This computational system can be described both as a neural network and algebraically.

#### The neural network and the algebraic representation

The two representations are shown side-by-side in Fig. 2. From a neurobiological perspective, the computational system is a neural network with an input level, an intermediate level with recurrent connections, and an output level (see Fig. 2A). These recurrent connections agree with the fact that, in the CNS, neuronal groups are connected both in a feedforward (away from the sensory input) and a feedback (towards the sensory input) direction. As is common in neuroscience, I assume that the CNS learns optimal weights for the neural network’s connections [28, 42, 43]. Calculating these optimal weights is a computational challenge and is often considered a part of machine learning. Here, I follow a different approach: I establish a correspondence between the neural network and an equivalent algebraic approach for which an optimal solution is known. I use an LQG optimality criterion that quantifies the objective to stay upright using control signals that stay within the torque limits of the mechanical system [9]. The LQG-optimal solution computes (1) an optimal internal state estimate 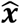 by integrating a linear differential equation (the Kalman filter) that takes as input an efference copy ***u*** and sensory feedback ***y***, and (2) an optimal control action ***u*** by multiplying the state estimate 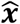 by the LQR gain *-K* (see Fig. 2B).

**Fig. 2:**
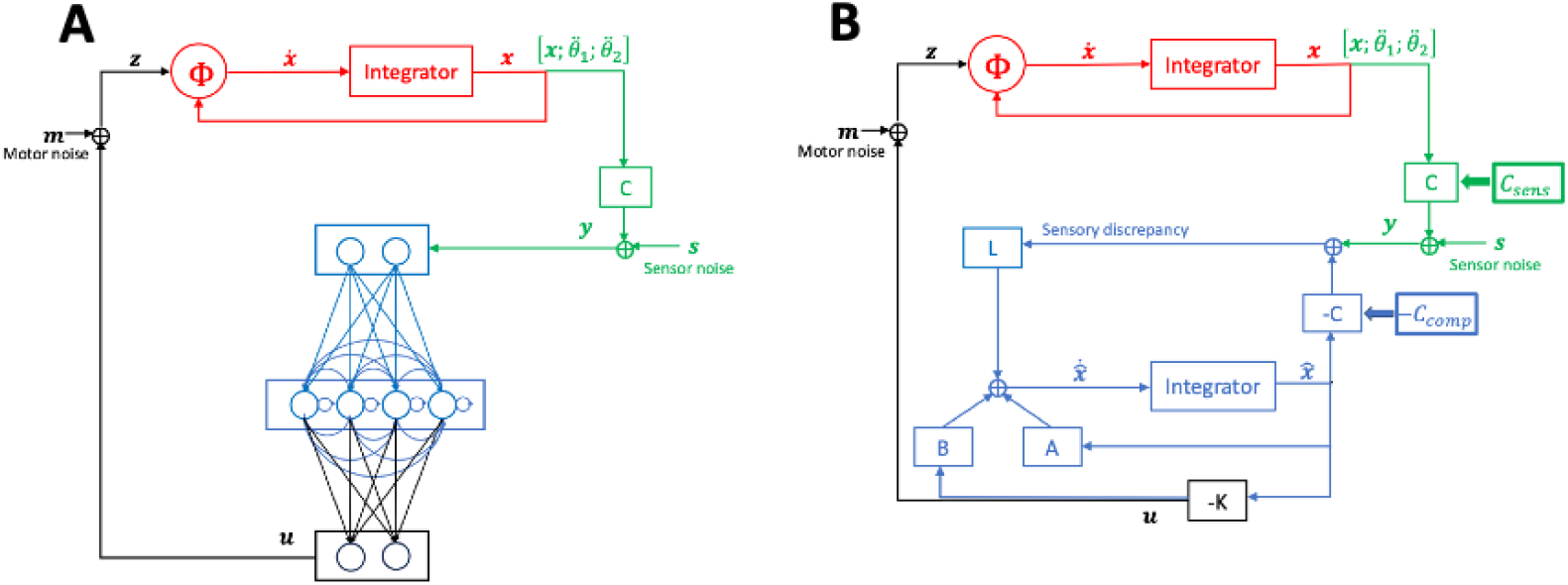
Closed-loop feedback control model without SMD. (A) Neural network representation (see text). (B) Corresponding algebraic representation (see text).

The computational system is based on (1) a linear forward model 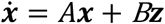 that approximates the nonlinear mechanical system 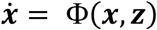 , and (2) a linear sensory model ***y*** = *C*_*Comp*_***x*** + ***s*** of the input to the computational system. I assume the matrices *A, B* and *C*_*Comp*_ to be known/learned and therefore can compute the LQG-optimal values for *L* (the Kalman gain) and -*K* (the LQR gain). It follows that the optimal recurrent weights are *A* − *BK* − *LC*_*Comp*_. To prevent possible misunderstandings, the equivalence between the neural network in Fig. 2A and the algebraic formalism in Fig. 2B is only used to demonstrate its neural plausibility; no neural network implementation with nonlinear properties has been built.

The computational system receives input from a sensory system, and it implements an internal model for this system. In Fig. 2B, the sensory system is characterized by the matrix *C*_*Sens*_ and its internal model by the matrix *C*_*Comp*_. I consider a sensory system for muscle spindle proprioceptive feedback, which is characterized by the following properties: (1) the firing rates of the muscle spindle primary afferents scale with intrafusal fiber acceleration [38-41], and (2) this intrafusal fiber acceleration is controlled by fusimotor neurons that cancel the reafferent acceleration component [10, 29-31] resulting in exafferent acceleration feedback. To avoid clutter, the fusimotor control of the sensory system is not included in Fig. 1 (panels B and C) and Fig. 2, but it is included in the corresponding Fig. 1 in [10]. The matrix *C*_*Sens*_ linearly combines total acceleration feedback (exafferent plus reafferent) and fusimotor input, which cancels the reafferent acceleration component, resulting in exafferent acceleration feedback [10].

The exafferent acceleration feedback of the DCIP is a nonlinear function of the state variables, but it can be approximated linearly at the unstable fixed point. The coefficient matrix of this linear approximation is the lower half of the matrix *A* in Eq. 2 and is denoted by *C*_*Comp*_. The resulting linear sensory internal model is not full rank, and therefore the state can only be estimated using a state observer (see Eq. 1). This model is described in more detail in [10] and in the Methods (see *A mechanical model for standing balance constrains the relation between exafferent joint acceleration and the state variables*).

#### The LQG solution

The LQG solution for a computational system without SMD compensation is the following linear dynamical system:

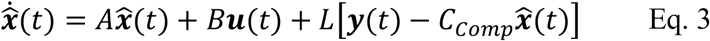

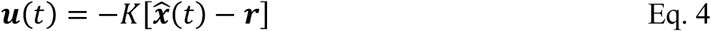

Eq. 3 is a state equation and is also called a state observer; if *L* is the Kalman gain, it is called the Kalman filter. The state observer specifies the derivative of the state estimate as the sum of two components: (1) an expected state derivative 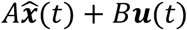 , and (2) a feedback-based correction 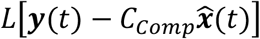. The state observer is a mathematical specification of the powerful idea that perception is the result of a process that combines what the CNS already believes (the current state estimate 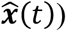) with sensory information ***y***(*t*) and a memory of its control signal ***u***(*t*).

Eq. 4 is an output equation, and it computes a control signal ***u***(*t*) by scaling the difference vector 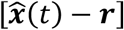 , in which ***r*** is the target state that specifies the movement objective. For balance control, ***r*** is the angle relative to gravity of the line that joins the center of gravity (CoG) with the area of support. People prefer a slightly leaned forward position of approximately 0.0156 radians [4], and therefore ***r*** = [0.0156,0.0156,0,0]^*t*^ is a plausible choice. For simplicity, the target state ***r*** was omitted from Fig. 1 and Fig. 2. In engineering terms, the combined state and output equation are a filter of the sensory feedback ***y***(*t*) and the target state ***r*** that produces the control signal ***u***(*t*) as its output.

The state observer in Eq. 3 is not needed if the sensory feedback is full-state and noise-free (i.e., ***y***(*t*) = ***x***(*t*)). In that case, in the output equation Eq. 4, 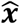 is replaced by ***x***. If the sensory feedback is full-state but noisy, the state observer (with *C*_*Comp*_ equal to the identity matrix) is used to optimally combine the noisy feedback with the forward model.

#### Expected versus predicted states

At this point, it is useful to make a sidestep to explain the difference between expected and predicted states. I do this because the expected state derivative 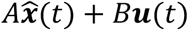 corresponds to a term in the discrete time state observer that has also been called a prediction [44-46], whereas in this paper, I use prediction in a different meaning. The discrete time state observer involves the following two-step operation:

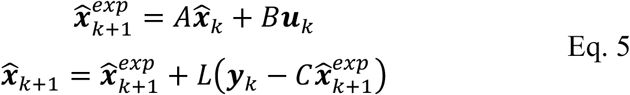

The time between consecutive integer-valued time points *k* and *k* + 1 is *δ*, and the relation between the continuous and the discrete time axis is given by *t* = *kδ*. I use 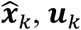 and ***y***_*k*_ as a shorthand for 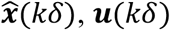 , ***u***(*kδ*) and ***y***(*kδ*). For notational simplicity, I have reused the symbols *A, B, L* and *C* of the continuous time state observer in Eq. 3, but the first three refer to different matrices than the corresponding ones for the continuous time observer. I will return to this when describing Eq. 15.

The relevant term in Eq. 5 is 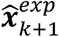 , which I will denote as the expected state, but in other papers it is also denoted as the predicted state or the a priori state estimate. The expected state 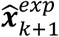 predicts the state estimate at time (*k* + 1)*δ* from the state estimate at time *kδ* (i.e., 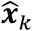) and the efference copy ***u***_*k*_. Crucially, this is not a prediction over a neurobiologically meaningful interval, but over a discretization interval of length *δ* that is chosen small enough to study the properties of the continuous state estimate 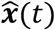 in discrete time. For prediction over a neurobiologically meaningful interval, I use *d* = *SMD/δ* to denote the SMD length as an integer number of intervals of length *δ*. In a later section (*The convolution predictor*), I will motivate a prediction of the state at time (*k* + *d*)*δ* = *k* + *SMD* from a state estimate at time *kδ* and the efference copies ***u***_*k*_, ***u***_*k*+1_, … , ***u***_*k*+*d*_. This is very different from the state observer, which combines sensory feedback and the most recent efference copy, but does not meaningfully predict beyond the time of the sensory feedback; its discrete time version only predicts over an arbitrary discretization interval.

### Gain scaling

Gain scaling is depicted in Fig. 1B. Gain scaling modifies the output equation Eq. 4. The output of this equation scales with the gain −*K*, and the larger its magnitude |*K*|, the more aggressive the corrections that are intended to bring the mechanical state to its target. With increasing SMD, it becomes more likely that corrections destabilize the mechanical system. This is because the stability of a closed loop system depends on the gain of the open loop system (the mechanical and the sensory system, followed by the observer and the controller −*K*) at a critical frequency, the frequency for which the open loop transfer function flips the phase by 180 degrees. To achieve stability, at this critical frequency, the amplitude of the open loop output gain must be less than 1. Crucially, this critical frequency decreases with the SMD and brings it in a range where the open loop output gain typically is higher. Now, by downscaling |*K*|, the open loop output gain decreases, ideally below 1 at the critical frequency.

For a mechanical system, the output gain −*K* is a matrix and, in principle, every matrix element could be scaled separately. This is also the scaling scenario depicted in Fig. 1B, and optimal scaling in this scenario would thus be equivalent to knowing the optimal output gain for a nonzero SMD. Because there is no theory about optimal output gains for a nonzero SMD, this type of optimal scaling is computationally infeasible. In this paper, I only consider optimal global downscaling of the output gain: replace −*K* (the LQR gain for SMD=0) by −*αK*, in which *α* optimizes the LQR objective function.

### The convolution predictor

The convolution predictor computes a control signal by applying an output gain to a prediction of the future state ***x***(*t* + *SMD*). In its simplest form, the convolution predictor is obtained as the solution of a delayed-input forward model, and this is also how it was introduced under the label finite spectrum assignment (FSA) [11-17]. Here, I take the same starting point but deviate from the existing literature by estimating the initial condition of the solution by a state observer that is based on the delayed-input forward model. This generalizes the method to situations with noisy and incomplete sensory feedback, and it does so optimally if the state observer uses the Kalman gain.

#### Solving the delayed-input forward model

The delayed-input forward model describes the dynamics of the mechanical system as follows:

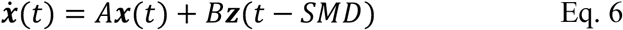

In this forward model, there is a delay of SMD seconds between the noisy control signal ***z*** (= ***u*** + ***m***) and the time at which it arrives in the mechanical system. This delay reflects the fact that the control signal is based on sensory feedback that is generated SMD seconds before its arrival in the mechanical system.

The future state ***x***(*t* + *SMD*) is obtained by evaluating the solution of Eq. 6 at time *t* + *SMD*. This solution can be found in the specialized literature [13, 17] or derived from more general results in a textbook [47]:

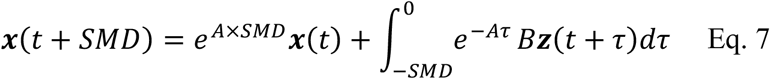

in which the multiplication sign (×) is only added for readability. The ingredients for this solution are the initial condition ***x***(*t*) and the noisy control signals ***z***(*t* + *τ*) for −*SMD* < *τ* < 0. From Eq. 7, one can obtain an estimate of the future state ***x***(*t* + *SMD*) by replacing the initial condition and the noisy control signals by estimates. This is straightforward for the noisy control signals, which can be replaced by the efference copies ***u***(*t* + *τ*). Crucially, this operation involves the assumption that the CNS has access to a memory buffer that stores the efference copies for the most recent interval of SMD seconds.

The initial condition can be estimated from the sensory feedback ***y***(*t*), and this will be described in the next section, *The delayed-input state observer*. There, it will be described how, from the sensory feedback ***y***(*t*) one can obtain an estimate of ***x***(*t* − *SD*), the state of the mechanical system at the time the sensory feedback was generated. With the estimate 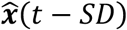 as the initial condition, application of Eq. 7 results in an estimate of ***x***(*t* + *MD*), the state of the mechanical system SMD seconds after the sensory feedback was generated. I will denote this estimate by 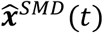:

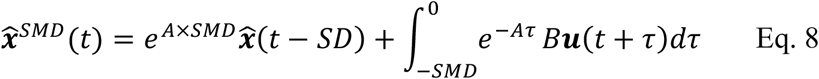

A potentially confusing aspect of this equation is that it combines quantities that are expressed on different times axes: 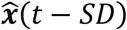 estimates a state at time *t* − *SD* on the time axis of the mechanical system, but this is computed at time *t* on the time axis of the computational system (CNS), which is also used to express the efference copies ***u***(*t* + *τ*). In other words, at time *t* on the time axis of the computational system, a prediction 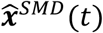 is computed that estimates ***x***(*t* + *MD*) on the time axis of the mechanical system, and for that it uses efference copies ***u***(*t* + *τ*) on the computational system’s time axis.

Note that the convolution predictor does not have to know the separate sensory and motor delay (SD, resp., MD), but only their sum SMD. This is because it uses an initial condition 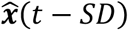 of which it doesn’t have to know the implied time axis; it only must know the total time span of the memory buffer of efference copies.

#### The delayed-input state observer

The delayed-input state observer computes the initial condition 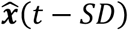 that is used in the convolution predictor in Eq. 8. This state observer is based on the delayed-input forward model in Eq. 6, and is written as follows:

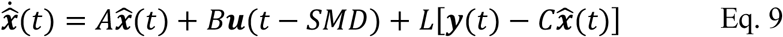

The difference with the state observer in Eq. 3 is the expected state derivative 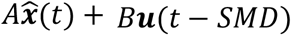 , which is taken from the delayed-input forward model instead of the forward model with non-delayed input. If the matrix *L* in Eq. 9 is the Kalman gain, then the delayed-input state observer produces an optimal estimate of ***x***(*t* − *SD*), namely the estimate with the smallest mean-squared error. This follows from the following facts: (1) the sensory input ***y***(*t*) is a noisy version of ***Cx***(*t* − *SD*), and (2) at the time the sensory input is generated, the systematic (noise-free) part of the mechanical system input is ***u***(*t* − *SMD*). This may be confusing because ***x***(*t* − *SD*) and ***u***(*t* − *SMD*) are expressed on different times axes: ***x***(*t* − *SD*) is expressed on the time axis of the mechanical system and ***u***(*t* − *SMD*) on the time axis of the computational system, where control signals are generated MD seconds before they arrive in the mechanical system.

The delayed-input state observer requires that the CNS has access to a memory buffer that stores efference copies ***u***(*t*) of the control signals that were produced in the most recent SMD interval (i.e., between *t* − SMD and *t* seconds). It uses the oldest efference copy ***u***(*t* − *SMD*) from this buffer to evaluate the state observer in Eq. 9.

#### Approximate convolution

As will be explained in the following, it is likely that the convolution operation in Eq. 7 is a computational challenge for the CNS. Therefore, it is useful to investigate whether approximate convolution could be as effective for SMD compensation as the full-blown convolution operation. To introduce different approximate convolution schemes, it is useful to start from the discrete time version of Eq. 8. In the derivation of this discrete time version, I assume that ***u***(*t*) is a piecewise constant function, with piece length equal to *δ* sec., the defining time step for the discrete time version. The discrete time quantities that correspond to the predictions 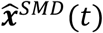 , the state estimates 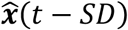 , and the efference copies ***u***(*t*) are denoted by, resp., 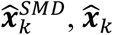 and ***u***_*k*_, with index *k* = *t*/*δ*. (The index *k* under the state estimate 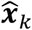 denotes the time at which the estimate is computed, and not the time at which the corresponding sensory feedback was generated.) Under the assumption of a piecewise constant ***u***(*t*), the discrete time approximation of Eq. 8 can be written as follows^1^:

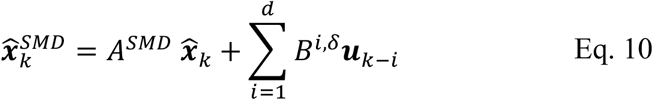

In this equation, *A*^*SMD*^ denotes the matrix exponential *e*^*A*×*SMD*^, *d* equals *SMD*/*δ*, and the efference copy weight matrices *B*^*i*,*δ*^ denote an integral that has an analytical solution:

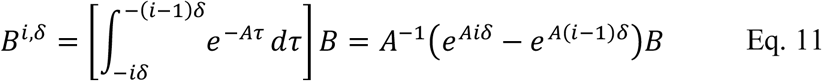

The convolution in the second term on the right of Eq. 10 is computationally heavy because it does not allow to reuse parts of previous computations. It is unknown whether the CNS has the required neural circuitry to successfully complete this job. In addition, this term involves a set of efference copy weight matrices *B*^*i*,*δ*^ that must be learned from experience with the mechanical system. Importantly, for an unstable mechanical system, the oldest efference copies are the most important ones, because the weights *A*^−1^(*e*^*Aiδ*^ − *e*^*A*(*i*−1)*δ*^)*B* increase exponentially with the buffer time index *i*. This exponential increase follows from the fact that an unstable system has a system matrix *A* with at least one positive eigenvalue. Thus, the largest weights are the most difficult to learn because they depend on the oldest memories, and these are likely to be the least accurate.

To investigate the consequences of different computational limitations of the CNS, I consider four predictors of different levels of complexity: (1) from passive dynamics only (i.e., using only the first term on the right side of Eq. 11), (2) under the assumption that the efference copy remains constant at the most recent value (i.e., using ***u***_*k*−1_ with a single weight 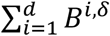 , (3) using the same average weight 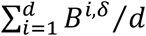 for all efference copies, and (4) using the full-blown mechanically motivated Eq. 10. The first two predictions do not make use of a memory buffer and therefore can be considered as extrapolations based on, respectively, only the current state estimate or the current state estimate plus the current control signal. The third predictor is a simple boxcar filter but requires a full memory buffer.

### The Smith predictor

Just like the convolution predictor, the Smith predictor computes a control signal by applying the output gain to a prediction of a future state. The Smith predictor assumes that the sensory feedback ***y***(*t*) is full-state and noise-free: ***y***(*t*) = ***x***(*t* − *SMD*). Now, if the SMD were 0, then ***y***(*t*) = ***x***(*t*) and the control signal would be obtained by multiplying ***y***(*t*) by the output gain: ***u***(*t*) = −*K****y***(*t*). (For simplicity, I ignore the target state ***r*;** see Eq. 4.) An easy way to introduce the Smith predictor is as a correction of the sensory feedback ***y***(*t*) for the unwanted component that is due to a nonzero SMD. This unwanted component is ***x***(*t* − *SMD*) − ***x***(*t*), the actual minus the desired sensory feedback.

The Smith predictor estimates the unwanted component using the forward model with non-delayed and noise-free input:

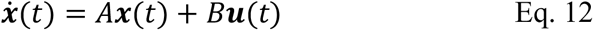

This estimate is the difference 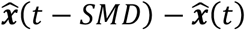 , in which 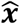 denotes a state estimate that is obtained from the solution of the forward model in Eq. 12. Subtracting this correction term from ***y***(*t*), the Smith predictor becomes 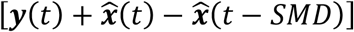 , which is then multiplied by the output gain to obtain the control signal. An alternative way to introduce the Smith predictor is to consider 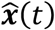 as an estimate of the current state and 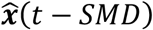 as a correction term for ***y***(*t*).

The solution of the forward model with non-delayed input (see Eq. 12) has a similar structure as the solution of the forward model with delayed input that was used for the convolution predictor (see Eq. 7):

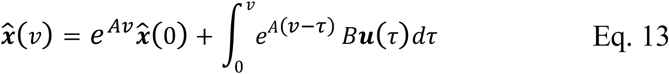

in which *v* can be *t* or *t* − *SMD*. There are several important differences with the convolution predictor, and these are best highlighted using the discrete-time version of Eq. 13. As for the convolution predictor, I assume that ***u***(*t*) is a piecewise constant function, with piece length equal to *δ* sec. The discrete time version of Eq. 13 can then be written as follows:

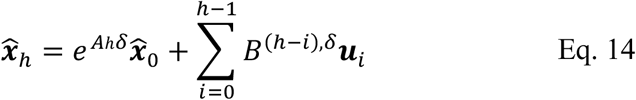

in which *h* can be *k* or *k* − *d*, and *B*^(*h*−*i*),*δ*^ is defined in Eq. 11. The first difference with the convolution predictor in Eq. 10 is that the initial condition 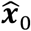 in Eq. 14 must be set once at the start of the time series and independently of the sensory feedback. The second difference is that the summation in Eq. 14 is not a convolution but an ordinary sum with number of terms increasing with *h*. This allows for a computationally efficient implementation based on this recursion:

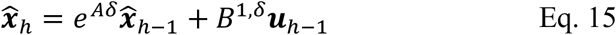

This recursion can be derived by inserting the definition of *B*^(*h*−*i*),*δ*^ (see Eq. 11) in Eq. 14. Eq. 15 is the discrete time forward model with non-delayed input and is identical to the first step in Eq. 5, but now with the formally correct system matrix *e*^*Aδ*^ and input matrix *B*^1,*δ*^; for simplicity, these matrices were denoted by *A* and *B* in Eq. 5. And the third difference is that a computationally efficient implementation of the Smith predictor requires a memory buffer of state estimates 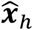 instead of efference copies ***u***_*h*_. To compute the correction term in the discrete time Smith predictor 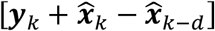 , this memory buffer must contain *d* elements, just like the memory buffer for the convolution predictor.

In sum, the Smith predictor estimates a future state by correcting full-state delayed sensory feedback. This correction is based on the solution of a forward model with non-delayed feedback, and it can be implemented in a computationally efficient way using a memory buffer of state estimates.

### The Smith predictor as a model for CNS computations

In motor control, the Smith predictor is far more popular than the convolution predictor, which is mostly discussed in the control theory literature [for an exception, see 16]. A probable reason for this is that the Smith predictor has played such an important role in the popularization of the concept of a forward model [19, 24, 25, 48]. However, the Smith predictor is not a useful model for CNS computations and this is for several reasons: (1) it cannot control an unstable mechanical system, (2) it ignores the fact that a plausible forward model must have a delayed input, whose learning and use requires a memory buffer of efference copies, and (3) it cannot be used in combination with a state observer.

First, the control theory literature has demonstrated that, under conditions that make it an interesting model for the CNS, the Smith predictor cannot control an unstable mechanical system [11, 20, 49]. The conditions that make the Smith predictor an interesting model for the CNS pertain to how well the forward model approximates the mechanical system. Specifically, to be an interesting model for the CNS, the Smith predictor’s performance (1) must be robust to some degree of approximation, and (2) must improve as the approximation becomes better. Although the control theory literature is clear in this respect, its results must be formulated carefully: there are examples of an unstable system that is controlled by the Smith predictor, but these do not involve a forward model that approximates the mechanical system. Instead, they involve a stable forward model that is totally unrelated to the unstable mechanical system [20].

Second, the Smith predictor ignores the fact that a plausible forward model must have a delayed input. The CNS can learn a forward and a sensory model only by associating motor signals (the input to the mechanical system) with sensory feedback (the signal that informs CNS about the mechanical system’s output). Because of the SMD, this learning-by-association is only possible if the CNS learns a forward model with delayed input, and this in turn requires that the CNS has access to a memory buffer of efference copies.

Third, when used as a model for CNS computations, the Smith predictor cannot be used in combination with a state observer. From an engineering point of view, it is straightforward to replace the sensory feedback ***y***(*t*) by the output of the delayed-input state observer in Eq. 9, but from the point of view of its possible neurobiological implementation, such a hybrid mechanism would be implausible. This is for two reasons. First, it would require that the CNS implements both the forward model with delayed input (for the delayed-input state observer) and the forward model with non-delayed input (for the correction term). Second, it would require that the CNS implements two memory buffers: one memory buffer that stores efference copies (for the delayed-input state observer), and another one that stores state estimates (for the correction term).

### Evaluating the performance of the SMD compensation mechanisms using simulation

I now give a general description of a set of the simulation studies with which I have evaluated the performance of the different SMD compensation mechanisms; the details are given in the Methods. The parameters of the two mechanical systems (a standing person and a rider-bicycle combination) were assigned realistic values, most of which were based on measurements (see Methods).

The model for the standing person (the DCIP) is nonlinear, and the one for the rider-bicycle combination is linear. Because the SMD compensation mechanisms depend on linear internal models, control of the nonlinear standing person system requires internal models that only approximate this mechanical system. In the simulations, these internal models were obtained as linear approximations of the mechanical system’s nonlinear EoM. In the control of the linear rider-bicycle system the internal models were identical to this system’s linear EoM.

I ran simulations using two sensory systems: (1) a neurophysiologically plausible sensory system that produces noisy exafferent acceleration feedback, and (2) a neurophysiologically implausible sensory system that produces noisy full-state feedback. In my report, I focus on the simulations with noisy exafferent acceleration feedback. If these simulations reveal an interesting pattern, I investigate their generality by checking whether this pattern is also present in simulations with noisy full-state feedback.

The time-varying input to the closed-loop control system involves sensor, motor and fusimotor noise. The noise amplitudes were set such that, without SMD, the root-mean-square deviation (RMSD) of the simulated CoG lean angle was equal to the RMSD of empirically observed CoG lean angles. The empirically observed RMSD for standing balance control (0.25 degrees) was taken from Figure 6 in [50], and the one for bicycle balance control (1.3321 degrees) was computed from a pilot dataset obtained from a study using a bicycle simulator [51]. The amplitude of the sensor noise was set equal to the amplitude of the combined sensory consequences of the motor and fusimotor noise. The motor and the fusimotor noise had identical amplitudes and their shared variance was 50%. These noise amplitudes were set separately for the exafferent acceleration and the full-state feedback; they were more than an order of magnitude larger for the full-state feedback.

The LQR gain −*K* minimizes an objective function that quantifies a double objective with a precision and an energetic component: stay close to the target position (precision) with as little effort as possible (energetic). I computed −*K* using parameters that weight the contribution of the precision and the energetic component equally, as quantified by realistic maximum values for the states and the joint torques. The Kalman gain *L* depends on the motor, fusimotor and sensor noise amplitudes, and I used the same values as for the noise signals that were fed into the mechanical and the sensory system.

Gain scaling starts from the LQR-optimal gain matrix −*K* for a non-delayed system. For every SMD, I computed the scalar scaling parameter *α* that minimizes the LQR objective function for a delayed system controlled using the output gain −*αK*. Depending on the SMD, it could happen that no *α* parameter value could be found that controls the mechanical system (i.e., the simulated standing person or rider-bicycle combination fell on the ground). In that case, it was concluded that gain scaling could not control the mechanical system for this SMD and larger ones.

The convolution predictor 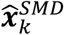 (see Eq. 10), was implemented in four different versions: (1) extrapolation based on the state estimate (passive dynamics only; without the second term on the right of Eq. 10), (2) extrapolation based on the state estimate plus the most recent efference copy (***u***_*k*−*i*_ = ***u****k*−_1_), (3) using the average efference copy weight 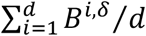 instead of *B*^*i*,*δ*^ (i.e., a boxcar filter), and (4) exactly according to Eq. 10.

The evaluation of the SMD compensation mechanisms is based on the largest SMD for which the RMSD CoG lean angle stays below an empirically determined maximum. For standing balance control, this maximum (1.5 degrees) was determined as the maximum RMSD that people tolerate and was taken from Fig. 6 (upper right panel) in [50]. For bicycle balance control, this maximum (15.11 degrees) corresponds to a turn radius that most people are not willing to take at the simulated speed (15.5 km/h) [10].

### Standing balance control

Fig. 3 shows the standing balance simulation results for four of the six SMD compensation mechanisms. At large SMDs, the simulated angular position may exceed 90 degrees, indicating that the person falls on the floor. Simulated falls happened for all SMD compensation mechanisms but, depending on the mechanism, this started at a different SMD. If a simulated fall happened, no results are shown for the corresponding SMDs in Fig. 3.

**Fig. 3:**
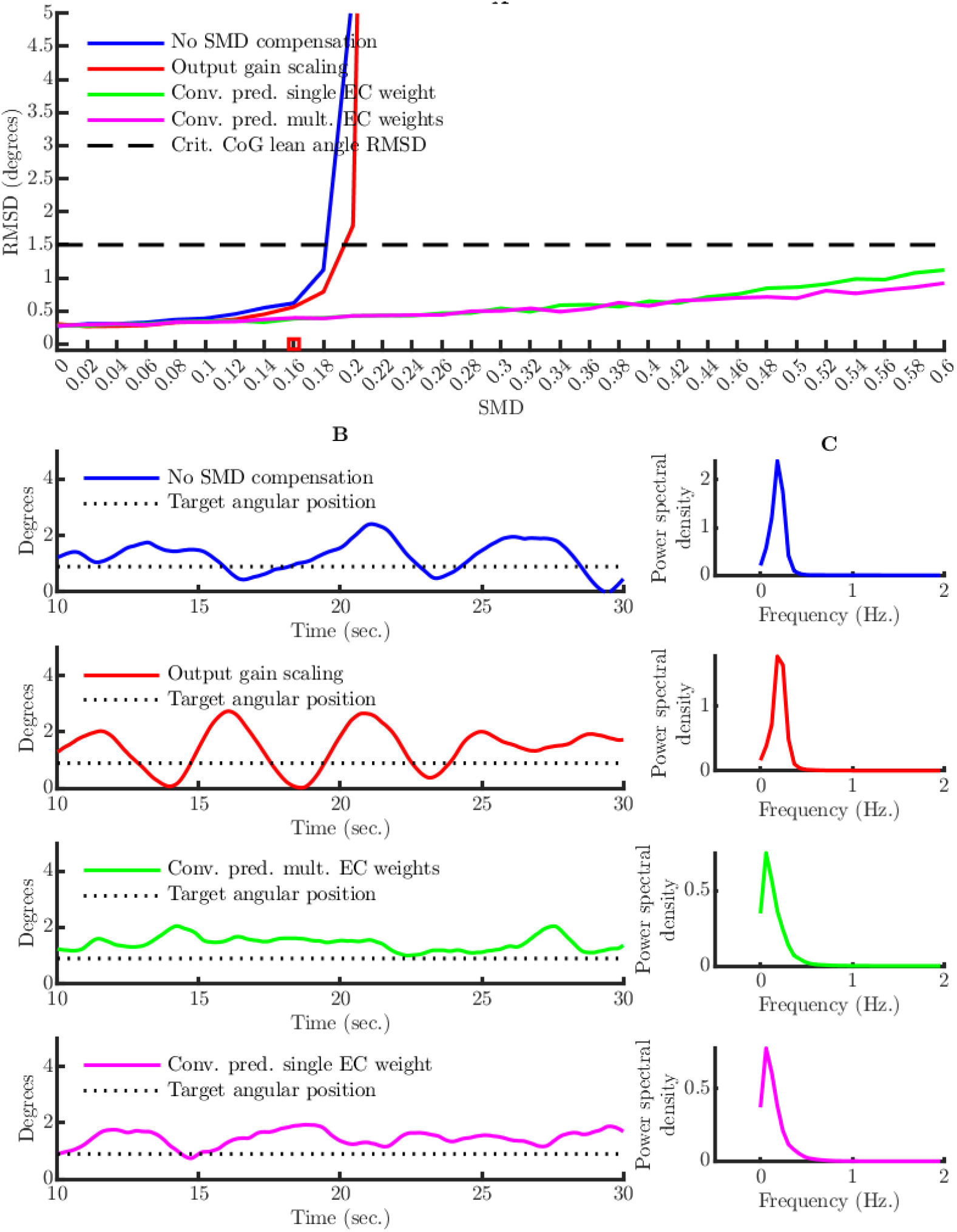
Standing balance simulation study results. (A) Average RMSD CoG lean angle for four SMD compensation mechanisms (including no SMD compensation), as a function of SMD. The horizontal dashed line indicates the maximum angular position RMSD that people tolerate and is based on empirical work [50]. (B) Example angular position time series for the SMD that is indicated by a red box in panel A (SMD=0.16 sec.). (C) Power spectral density of the angular position for the same SMD as in B.

Without a SMD compensation mechanism, the critical CoG lean angle RMSD was exceeded for SMD>0.181 sec. (see Fig. 3A). This SMD is slightly above the range of the neurobiological SMDs that have been reported in the literature: 0.1-0.16 sec. in [4, 52] and 0.14 sec. in [23]. As expected, output gain scaling produced smaller CoG lean angle RMSDs for all SMDs, and the critical RMSD was exceeded for a slightly larger SMD of 0.195 sec. (see Fig. 3A). The acceptable performance under no SMD compensation and output gain scaling depends on joint stiffness and damping (both at the ankle and the hip): lowering stiffness and damping below the measured values lowered the SMD that could be tolerated without exceeding the critical RMSD. When ankle joint stiffness and damping were set to zero, the maximum SMD that could be tolerated was only 0.05 sec.

The performance of the convolution predictor strongly depends on whether a full memory buffer of efference copies was used. The performance of extrapolation was so poor that the lines are not even visible in Fig. 3A: for passive dynamics only, the critical RMSD was exceeded for SMD>0.022 sec., and if also the most recent efference copy was used, the critical RMSD was exceeded for SMD>0.013 sec.

The versions of the convolution predictor that make use of a full memory buffer of efference copies have an excellent performance: both versions (with single and multiple efference copy weights) exceed the critical CoG lean angle RMSD only for a SMD larger than 0.6 sec. (see Fig. 3A). The precise maximum values are not shown in Fig. 3A; they are equal to 0.71 and 0.91 for, respectively, the single and the multiple weights version. It is useful to compare these values with the results of an empirical study in which the SMD was artificially increased by means of a robotic device [4]. In this study, humans could successfully compensate for an artificial SMD of 0.56 sec., well below the performance limit of a convolution predictor that makes use of a memory buffer of efference copies.

The performance of the convolution predictor depends on joint stiffness and damping: lowering stiffness and damping below the measured values lowered the SMD that could be tolerated without exceeding the critical RMSD. However, this effect was much less extreme than for the scenarios without SMD compensation and with output gain scaling. When ankle joint stiffness and damping were set to zero, the maximum SMDs that could be tolerated were 0.52 and 0.65 sec. for, respectively, the version with a single and multiple efference copy weights.

Two important results are (1) that the convolution predictor is only effective if it makes use of a full memory buffer of efference copies to predict a future state, and (2) that there is little difference between a computationally hard convolution using multiple efference copy weights and a computationally easy convolution using only a single weight. These results (necessity of a full memory buffer and sufficiency of a single weight) are not specific for noisy incomplete exafferent acceleration feedback; they were also obtained with noisy full-state feedback. Using the latter feedback, for extrapolation based on passive dynamics only and for extrapolation using the most recent efference copy, the critical RMSD was exceeded for, respectively, SMD>0.035 sec. and SMD>0.012 sec. And for the single and the multiple weights version of the convolution predictor, the critical RMSD was exceeded for, respectively, SMD>0.61 sec. and SMD>0.80 sec. Keep in mind that the noise level was set higher for full-state than for exafferent acceleration feedback to achieve an identical RMSD of 0.25 degrees at zero SMD for both feedback types.

I also confirmed the mathematical results for control performance of the Smith predictor when applied to an unstable mechanical system: even for the smallest SMD (0.001 sec.), the Smith predictor produced control signals that resulted in diverging CoG lean angles.

To give a more concrete picture of the simulated kinematics, in Fig. 3B, I show examples of CoG lean angle time courses for the effective SMD compensation mechanisms. Note the rhythmic pattern around 0.2 Hz for the scenario’s without SMD compensation and with output gain scaling. Also note that the average CoG lean angle exceeds its target value, which is due to the positive gravitational torque that acts on the body when it is in a slightly leaned forward position.

I computed the power spectral densities of the CoG lean angle time courses after removing the mean CoG lean angle. These power spectral densities all show the drop-off between 0 and 2 Hz that is observed in empirical CoG lean angle power spectral densities (see Fig. 3C) [53].

### Bicycle balance control

#### A mechanical model for bicycle balance control

There are two important differences between the physics of standing and bicycle balance control: (1) a bicycle’s area of support (AoS) is a line instead of a surface, and (2) balance control of a moving bicycle involves not only the gravitational but also the centrifugal force. A stationary bicycle is balanced when the combined CoG of rider and bicycle is above the line that connects the contact points of the two wheels with the road surface, the so-called line of support (LoS). Because of disturbances, this CoG cannot be exactly above this one-dimensional LoS for some time. Therefore, a bicycle is considered balanced if the CoG fluctuates around the LoS within a limited range, small enough to prevent the bicycle from skidding and/or touching the road surface.

On a moving bicycle, not only gravity, but also the centrifugal force acts on the CoG. Crucially, the centrifugal force is under the rider’s control via the turn radius [54]. The balance of a moving bicycle depends on the resultant of the gravitational and the centrifugal force: a bicycle is balanced if the direction of this resultant force fluctuates around the LoS without diverging. This dependence on the resultant force is also observed in walking, running, and skiing. Besides the forces that act on the CoG, there are also forces that turn the bicycle’s front frame, and some of these forces are independent of the rider’s action [55].

These rider-independent forces are responsible for the bicycle’s self-stability within a limited speed range [55]. For the present study, I use the bicycle model in Fig. 4. This model consists of three rigid bodies: front frame, rear frame (which includes the rider’s lower body and will also be denoted as the lower body), and the rider’s upper body. The positions of these three bodies are specified by three angular variables: steering (*δ*), lower body (*θ*_1_), and upper body (*θ*_2_) angular position. The lower and upper body angular positions are relative to gravity.

**Fig 4:**
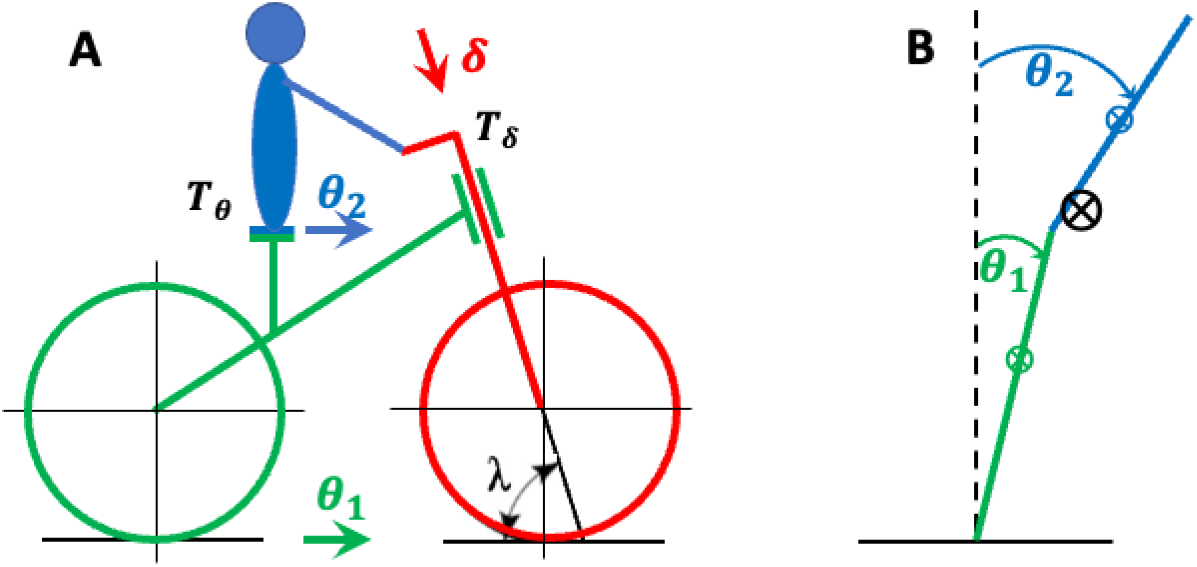
Kinematic variables of the bicycle model plus the rider-controlled forcing torques. (A) Side view. In green, the bicycle rear frame, characterized by its angular position *θ*_1_ over the roll axis (green arrow). In red, the bicycle front frame, characterized by its angular position *δ* over the steering axis (red arrow). In blue, the rider’s upper body, characterized by its angular position *θ*_2_ over the roll axis (blue arrow). In black, (1) the steering torque *T*_*δ*_ and the hip torque *T*_*θ*_, which are both applied by the rider, and (2) the steering axis angle *λ*. (B) Rear view. In green, the bicycle rear frame (including the lower body) angular position *θ*_1_. In blue, the rider’s upper body angular position *θ*_2_. The symbol ⨂ denotes the CoG of the lower body (in green), the upper body (in blue), and the combined CoG (in black).

Cycling involves a double balance problem, and until now I have only described the first problem, keeping the combined CoG of rider and bicycle above the LoS. The second balance problem pertains to the rider’s upper body only, and it involves keeping the upper body CoG above its own AoS, the saddle. I only consider balance over the roll axis (coinciding with the LoS), which corresponds to upper body movements to the left and the right. I thus ignore balance over the pitch axis (perpendicular to the LoS and gravity), which corresponds to upper body movements to the front and the back, typically caused by forward accelerations and braking.

For both balance problems (with respect to the combined and the upper body CoG), the relevant control action must produce a torque over the roll axis. Within the constraints of our kinematic model, there are two possible control actions: (1) turning the handlebars (using steering torque *T*_*δ*_), and (2) leaning the upper body (using hip torque *T*_*θ*_). At this point, it is convenient to make use of Fig. 4B, which shows the geometry of a DCIP but with different forces acting on it. By turning the handlebars, the contact point of the front tire (represented by the green rod) with the road surface moves to the left or the right, and this changes the position of the combined CoG relative to the LoS. In the bicycle reference frame (in which the LoS is one of the axes) this corresponds to a centrifugal torque in the direction opposite to the turn (a tipping out torque). Steering in the direction of the lean produces a tipping out torque that brings the combined CoG over the LoS. This is called steering to the lean/fall.

The second control action is leaning the upper body, which can bring the upper body CoG above the saddle in a direct way. This deals with the second (upper body specific) balance problem. However, leaning the upper body cannot deal with the first balance problem (bringing the combined CoG above the LoS) in a direct way (i.e., without turning the handlebars). A crucial argument in favor of this claim is that a bicycle with a locked steer cannot be balanced; not a single case has been reported. As an aside, it must be noted that leaning the upper body can deal with the first balance problem (the one with respect to the combined CoG) in an indirect way via the front frame: leaning the upper body to one side will make the front and the rear frame lean to the other side (by conservation of angular momentum) and, depending on the geometrical properties of the bicycle, this lean may turn the front frame to the same side [55, 56].

Starting from the kinematic bicycle model in Fig. 4, I formulated EoM for the mechanical system, and these are derived in the Methods. These EoM are an extension of a linear 2-DoF benchmark bicycle model that has been extensively validated [55]. The extension involved replacing the rear frame of the benchmark model by a linearized DCIP of which the lower joint is actuated by steering instead of by joint-crossing muscles [57]. This model is called the Benchmark Double Pendulum (BDP, see Methods).

#### What sensory information informs the CNS about the lean angle?

One of the most challenging aspects of bicycle balance control pertains to the sensory feedback that informs the CNS about the lean angle (combined CoG angular position). A more detailed description of this problem is given elsewhere [10], and here I only summarize it. The problem is best understood by assuming that a bicycle is balanced using the same type of proprioceptive feedback as a standing person: ankle and hip joint accelerations over the same axis as the angular position the person wants to control. Because the joint between the rear frame and the road surface is not actuated by joint-crossing muscles, it also does not provide proprioceptive feedback. Only the hip can be actuated by joint-crossing muscles and therefore only these muscles are a source of roll axis proprioceptive feedback. Not only does the CNS receive no proprioceptive feedback about the joint between the rear frame and the road surface, it also has no actuator to control this joint in a direct way. This joint can only be controlled indirectly by steering and rotating the hip joint. For this reason, the rider-bicycle combination belongs to the category of underactuated systems [58].

#### Bicycle balance simulation results

Fig. 5 shows simulation results for the four versions of the convolution predictor. It does not show the simulation results for the scenario without SMD compensation and the one with output gain scaling. The performance in these two scenarios is so poor that the lines would not be visible in the figure. For both scenarios, a 0.0025 sec. SMD resulted in lower body (bicycle rear frame) angular positions that were so large that the bicycle touched the road surface.

**Fig. 5:**
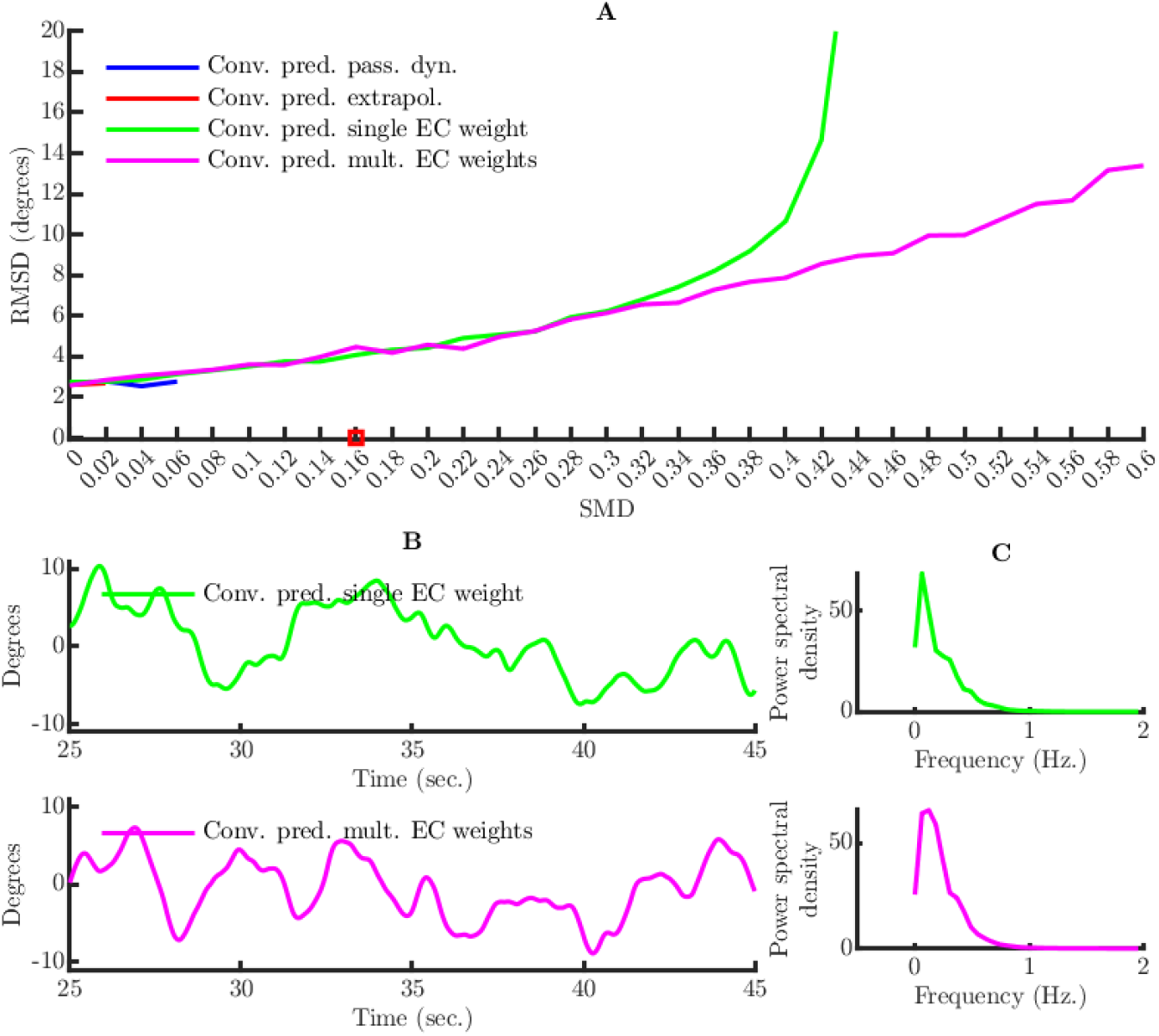
Bicycle balance simulation study results. (A) Average RMSD CoG lean angle for four SMD compensation mechanisms as a function of SMD. (B) Example angular position time series for the SMD that is indicated by a red box in panel A (SMD=0.16 sec.). (C) Power spectral density of the angular position for the same SMD as in B.

The performance of the convolution predictor strongly depends on whether a full memory buffer of efference copies was used. The performance of extrapolation was poor: for extrapolation based on passive dynamics only, the bicycle touched the road surface for SMD>0.07 sec., and for extrapolation using the most recent efference copy, it touched the road surface for SMD>0.035 sec. These critical SMDs are far below the neurobiological SMDs that are reported in the literature [4, 52, 59, 60].

A much better performance was obtained using the two versions of the convolution predictor that make use of a full memory buffer of efference copies (single and multiple efference copy weights). For the convolution predictor with a single efference copy weight, the bicycle started to touch the road surface for an SMD of approximately 0.41 sec. And for the convolution predictor with multiple efference copy weights this happened for an SMD of approximately 0.58 sec.

Different from standing balance control, for bicycle balance control, there is no experimentally determined critical lean angle RMSD beyond which cyclists feel too uncomfortable. I therefore used a rougher measure based on a plausible maximum for the combined CoG lean angle amplitudes. I calculated the lean angle amplitude that was required to take a 7 m. diameter corner at a speed of 15.5 km/h. (the speed used in the simulations) and found this to be 15 degrees inwards to the turn. Based on my own experience, this cornering task is feasible but challenging for experienced amateur mountain bikers. I assume that, without experience, this lean angle will feel uncomfortable, and I therefore calculated the SMD for which this critical lean angle amplitude was exceeded in more than one percent of the time. For the convolution predictor with a single efference copy weight, this happened for SMD>0.27 sec., and for the convolution predictor with multiple efference copy weights it happened for SMD>0.28 sec.

Thus, the convolution predictor is only effective if it makes use of a full memory buffer of efference copies to predict a future state, but there is little difference between a computationally hard convolution using multiple efference copy weights and a computationally easy convolution using only a single weight. These results (necessity of a full memory buffer and sufficiency of a single weight) are not specific to incomplete exafferent acceleration feedback because they were also obtained with noisy full-state feedback. With this noisy full-state feedback, the only SMD compensation mechanisms that are effective for a neurobiologically plausible SMD are the two versions of the convolution predictor that make use of a full memory buffer of efference copies.

As for standing balance control, I confirmed the mathematical results for control performance of the Smith predictor: even for the smallest SMD (0.001 sec.), the Smith predictor produced diverging upper and lower body angular positions.

For the two successful combinations, Fig. 5B and 5C show example combined CoG angular position time courses and their corresponding average psds. These are added for completeness but it is not informative to compare them to the angular position time courses when cycling on the public roads. In fact, the latter task involves road boundaries, which requires control actions (steering to stay between the road boundaries) that are unrelated to balance control.

## Discussion

I have described and evaluated three SMD compensation mechanisms: gain scaling, the convolution predictor, and the Smith predictor. These mechanisms are implemented using control theory results for linear dynamical systems [7, 8]. Dependence on results for linear dynamical systems can be motivated for balance control because the state of the body stays in a part of state space where a linear approximation of the EoM is accurate. I have evaluated these SMD compensation mechanisms both theoretically and by means of simulations. These are my conclusions:

1. The necessity of a SMD compensation mechanism depends on the mechanical system: no SMD compensation mechanism is required to balance a standing person with realistic joint stiffness and damping, but such a mechanism is required to balance a rider-bicycle combination. For conclusions 2-5, I focus on the rider-bicycle combination.
2. Compensating for a SMD requires the convolution predictor. This predictor involves a convolution over a memory buffer of efference copies.
3. The convolution predictor requires an initial condition obtained from a state observer that is based on a delayed-input forward model. Updating this state observer requires the oldest efference copy in the memory buffer of efference copies.
4. Compensating for a SMD does not require a computationally hard convolution with multiple efference copy weights; a similar performance is obtained with a computationally easy boxcar convolution that uses only a single weight.
5. Gain scaling is an effective SMD compensation mechanism but is not sufficient to compensate for a neurobiological SMD.
6. The Smith predictor is an ineffective and neurobiologically implausible SMD compensation mechanism for an unstable mechanical system.

### Limitations of this study

I now discuss three of the study’s main limitations: (1) the small set of mechanical systems and parameter configurations, (2) the absence of a systematic robustness evaluation, and (3) the confinement to a single task. First, it is difficult to rule out that the simulation results would have been different if different mechanical systems would have been used. Mechanical systems can differ in several aspects, such as geometry, mass distribution, joint stiffness and damping, and the type of control input (torque, impedance and stiffness modulation, etc.). With respect to geometry, for the rider-bicycle mechanical system, I could not obtain the EoM for a more realistic form of cycling and therefore considered a restricted form of cycling in which the rider keeps his legs still, does not use them to carry weight, and relies on a motor for propulsion. However, most cyclists transfer a large part of their weight to the pedals and the handlebars; downhill mountain bikers (the balance artists in the cycling community) even do this for the full 100 percent. For more realistic simulations, we need EoM for a rider-bicycle mechanical system in which (1) the lower body is no longer a part of the rear frame, and (2) the AoS for the combined upper and lower body is formed by saddle, pedals, and handlebars. Such a mechanical system also implies a very different sensory system for the exafferent acceleration feedback because the forces (gravitational and centrifugal) now produce exafferent acceleration feedback in both the upper and the lower body. If the rider’s legs carry weight (i.e., the saddle is at least partially unloaded) then the between-leg difference in load is informative about the resultant of the gravitational and the centrifugal force: because load is reflected in muscle tension, information about this resultant force can be obtained from leg muscle exafferent acceleration feedback.

Mechanical systems can also differ with respect to joint dynamics, and here I must distinguish between passive and active dynamics. Passive dynamics are all aspects of a mechanical system’s dynamics that do not depend on the system’s input, and it includes mass distribution, joint stiffness and damping. And active dynamics are all aspects of dynamics that depend on the input to the system, such as torque, impedance and stiffness. With respect to passive dynamics, I investigated the effect of mass distribution, joint stiffness, and joint damping on the simulation results, but not in a systematic way. In the Results, I reported on a few findings with respect to the stabilizing effect of joint stiffness and damping. With respect to active joint dynamics, the models only involved torque input, even though there are good arguments for the role of impedance and stiffness modulation [61, 62].

Second, this study is limited because it did not systematically investigate the robustness of the control performance to inaccuracies of the internal models. Robustness is relevant because the CNS must learn the internal models through experience with the mechanical system, and it is unrealistic to expect that these learned internal models exactly capture the mechanical system’s true dynamics (for the forward model) and the sensory feedback’s relation with the state variables (for the sensory model). This issue is especially relevant for LQG control with incomplete feedback because there is an example of an LQG controller with arbitrarily small stability margins [63], which makes it highly susceptible to model deviations.

Although I did not systematically investigate robustness to internal model inaccuracies, the simulations of standing balance control using the nonlinear DCIP do provide information about the robustness of the linear convolution predictor. In fact, the linear convolution predictor could control the nonlinear DCIP for SMDs that far exceed neurobiologically plausible SMDs.

Third, this study is limited because of its focus on balance control. Balance control is a form of continuous motor control without a beginning or end. This differs from pointing, reaching and grasping, which pertain to episodes with an obvious beginning and end. Such tasks naturally lead to a distinction between feedforward and feedback control, and this distinction is closely related to the SMD. Between the beginning of an action and SMD seconds later, no sensory feedback has arrived in the CNS, and therefore only feedforward control is possible; when the first sensory feedback arrives, the CNS can switch to feedback control. During this period of feedforward control, for a convolution predictor that integrates over the full memory buffer of efference copies (no extrapolation), the prediction changes as more efference copies are added to the memory buffer. I did not investigate whether such a mixed feedforward-feedback convolution predictor can model the movement trajectories in a pointing, reaching or grasping task.

### What is a plausible neurobiological implementation of the convolution predictor?

The Smith predictor has received a lot of attention in neuroscience because it has been linked to a neurobiological implementation with a central computational role for the cerebellum as a forward model with non-delayed input. This implementation involves two cortico-cerebellar loops [19]: one loop feeds a non-delayed state estimate 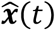 into the motor cortex, and the other loop estimates the delayed sensory feedback ***y***(*t*) by the delayed state estimate 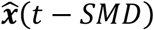.

Inspired by this neurobiological implementation of the Smith predictor, I now propose one for the convolution predictor (see Fig. 6). I hypothesize that the cerebellum computes convolutions over a memory buffer of efference copies. For the convolution over the full memory buffer (second term on the right side of Eq. 8), the convolution operator acts like a low-pass filter (see Fig. 6), and for the delayed-input state observer, the convolution operator selects the oldest efference copy from the memory buffer. The hypothesis that the cerebellum operates on a memory buffer of efference copies agrees with the fact that the cerebellum receives efference copies from the motor cortex, primarily through the cortico-ponto-cerebellar pathway. And the corollary that the cerebellum acts like a low-pass filter agrees with the fact that the most prominent symptom of cerebellar damage is a loss of smooth movements; cerebellar patients’ movements are erratic, uncoordinated, and incorrectly timed. This pattern is denoted as dysmetria.

**Fig. 6:**
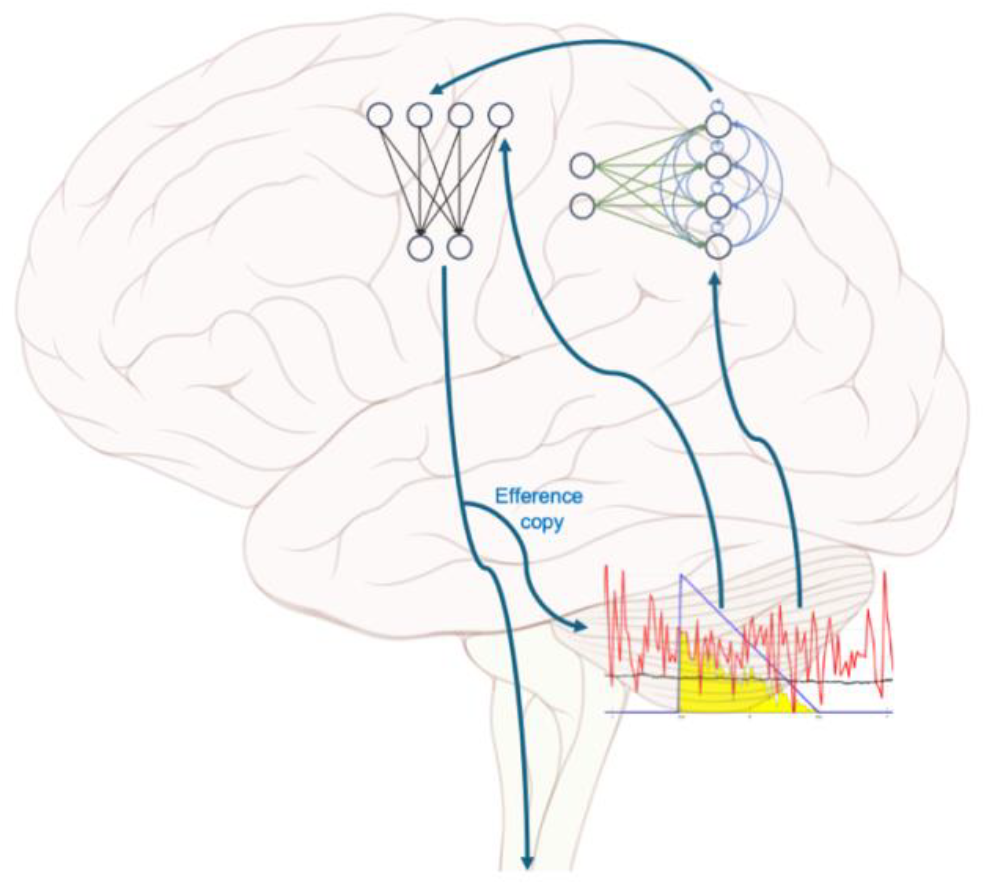
A possible neurobiological implementation of the convolution predictor. The cerebellum computes convolutions over a memory buffer of efference copies and sends output to the parietal and the motor cortex. A graphical representation of the convolution operation is superimposed on the cerebellum: the red signal is the efference copy input, the blue signal is the convolution kernel, and the black signal is the convolution output (the size of the yellow area under the thin grey line). The parietal cortex implements a state observer as a recurrent neural network and the motor cortex implements a feedforward neural network.

I hypothesize that the state observer is implemented in the parietal cortex. This agrees with the fact that the parietal cortex receives direct and short-range input from the sensory areas and possesses the recurrent connectivity necessary to implement a differential equation. Finally, I hypothesize that the motor cortex combines input from the parietal cortex (the state estimate) and the cerebellum (the outcome of the convolution over the memory buffer of efference copies) and computes a control signal using a feedforward neural network.

### Spinal mechanisms for motor control

The less the number of synapses between sensory input and motor output, the smaller the SMD. Therefore, it may be that SMD compensation mechanisms are less important if control mechanisms can be transferred to the spinal cord. To be plausible, spinal neural circuity must allow for computations that can realize the flexible motor control that humans are able to; simple mono-synaptic reflex arcs are almost certainly insufficient [64].

An elegant mechanism has been proposed that adds flexibility to the spinal stretch reflex: stretch-reflex sensitization via gamma motor neuron activity. Gamma motor neuron activity increases the intrafusal fiber tension, making muscle spindles more sensitive to stretch: a stretch that would normally trigger few or no action potentials in the sensory afferents, would now trigger many more after sensitization. Via the spinal reflex arc, these action potentials would activate alpha motor neurons and cause muscle contraction. This mechanism has been proposed in different variants, such as Merton’s servo hypothesis [65] and Feldman’s equilibrium point (lambda) hypothesis [66]. Different from stretch-reflex sensitization via intrafusal fiber tension, these variants state that gamma motor neurons specify a desired muscle length.

In its simplest form, the hypothesis that alpha motor neurons are activated indirectly via gamma motor neurons is unlikely to hold. An important reason is that it predicts a time delay between alpha and gamma motor neuron firing, and this time delay has not been observed. Instead, alpha and gamma motor neurons are typically activated almost simultaneously [67], a phenomenon known as alpha-gamma co-activation. Alpha-gamma co-activation is consistent with a sensory role for gamma motor neuron activation: cancellation of the reafferent acceleration feedback, as proposed in [10] and in line with [31].

Indirect activation of alpha motor neurons via stretch-reflex sensitization has been extensively criticized in the literature, and alpha-gamma coactivation is only one of the arguments in this debate [66, 68, 69]. Here, I add an argument that follows from the fact that a bicycle can only be balanced by turning the handlebars. If this control action would depend on stretch-reflex sensitization, this would have to involve reflex arcs that involve the arm muscles. However, contrary to the spindles in the axial muscles of the lower torso, the spindles in the arm muscles cannot detect a stretch over the same axis as the gravitational and centrifugal torque that act on axial muscles of the lower torso. Thus, because reflex arcs operate locally (muscle spindles and associated alpha motor neurons belong to muscles of the same joint), they cannot be responsible for control actions that involve muscle spindles and alpha motor neurons that belong to distant segments of the spinal cord.

### Prediction in perception and motor control

Prediction is a crucial concept in many subfields of neuroscience, and it is useful to distinguish between prediction in perception and prediction in motor control. I have argued that, for motor control in the presence of a SMD, prediction operates at the level of the output equation. This contrasts with the perception literature, where prediction is claimed to operate at the level of the state observer [70-74].

In the perception literature, prediction usually appears as a part of predictive coding, a theory that characterizes perception as an interaction between top-down (predictive) and bottom-up (sensory) influences [75-77]. This resembles the state observer which is a mathematical specification of the idea that perception is the result of a process that combines the CNS’s prior belief (top-down) with sensory information (bottom-up) and a memory of its control signals. It is therefore no surprise that the state observer has been described from the perspective of predictive coding [71, 73, 74].

Predictive coding is a hierarchical state estimation network, inspired by the known anatomical connectivity of the visual cortex [78, 79]. It proposes a hierarchical updating scheme that repeats itself in the different levels of the visual hierarchy, each of which computes a state estimate. For reference, this is the discrete time delayed-input state observer:

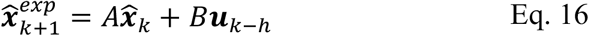

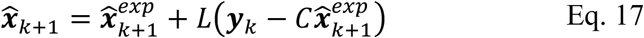

The relation between this state observer and predictive coding is not formally rigorous but rather loose, and this is in large part due to the numerous variants of predictive coding in the literature [80]. Here, I only consider the simplest variant, in which the efference copy ***u***_*k*−*h*_ is removed from Eq. 16. In more complex variants [81, 82], the efference copy is replaced by a so-called hidden cause, which represents the brain’s belief about the causes of the sensory input; the hidden states 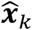 are an intermediate representation in between sensory input and hidden cause.

In all variants of predictive coding, ***y***_*k*_ represents the input from a lower to the next higher layer in the computational hierarchy. Thus, only in the first layer of the hierarchy, ***y***_*k*_ is a sensory signal with origin in the peripheral nervous system. In all higher layers, ***y***_*k*_ is a feedforward signal with origin in the CNS. The properties of this signal depend on the type of perceptual representation (w.r.t. tuning and receptive field size) in the sending layer.

The simplest form of predictive coding involves the following computational recipe:

1. A level in the predictive coding hierarchy sends a top-down prediction 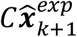 to the next lower level.
2. The prediction-receiving level also receives a bottom-up input ***y***_*k*_ from its next lower level.
3. The prediction-receiving level computes the sensory prediction error 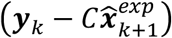 and sends it to the next higher (prediction-generating) level.
4. The prediction-generating level uses Eq. 17 to update the prediction.
5. After the update, the old prediction 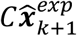 is assigned the value 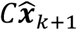 and the cycle repeats.

### Predictions and prediction errors for motor control

There are different views about the signal that the CNS sends to the spinal cord, and here I distinguish between a signal based on a prediction and one based on a prediction error. Starting with the first, this is the core claim of the present paper: the CNS sends a torque request that is based on the difference between a predicted and a target state: 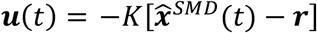. The alternative view has been proposed as a part of active inference [70, 83-85], which is an extension of predictive coding. In active inference, the CNS sends a sensory prediction error 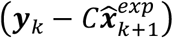 to the spinal cord. This prediction error is to be interpreted as a request for the spinal cord to minimize this error. Crucially, in the error signal 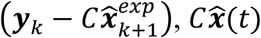 does not correspond to the predicted but to the desired sensory feedback; it has the same role as the target state ***r*** in the torque signal 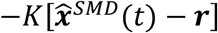. The prediction error 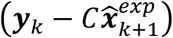 is thus a kinematic signal, and in that respect, it is like the input to a servo motor. However, it is different from a servo motor input in that the input signal must be minimized instead of realized.

In active inference, the alpha motor neurons are not controlled directly by the CNS but indirectly via reflex arcs that are assumed to minimize the sensory prediction error [70, 83-85]. The obvious candidate for this indirect activation is stretch-reflex sensitization via gamma motor neuron activity, which is also central in the equilibrium (lambda) point hypothesis [66]. An important difference between the two theories is that, under the equilibrium point hypothesis the kinematic input signal is realized by the reflex arcs, whereas under active inference it is minimized. However, many of the same criticisms that argue against indirect activation of alpha motor neurons via stretch-reflex sensitization [66, 68, 69] also apply to active inference. This includes the implausibility of bicycle balance control via stretch-reflex sensitization.

Whether based on predictions or prediction errors, to be useful, a computational mechanism must be able to control the states of the CNS-external mechanical system. That is, an important challenge lies in demonstrating that computations on CNS-internal states can control a CNS-external system. In this paper, I have demonstrated for two mechanical systems that this possible using a predictor that is computed as a convolution over a memory buffer of efference copies.

## Methods

### Matlab toolbox

All simulations were performed using the Matlab Balance Control (BalCon) toolbox that is shared in the supplementary information, together with the scripts that produced the simulation results, including the figures. For every mechanical system, the BalCon toolbox contains one function that computes the EoM and its linearization. The toolbox’s core functions are generic, in the sense that they can be used for all mechanical systems for which both the nonlinear and the linearlized EoM are provided, and for all sensory systems that can be linearized.

### A mechanical model for standing balance constrains the relation between exafferent joint acceleration and the state variables

Based on results from sensory neurophysiology, it can be argued that muscle spindle output is proportional to the exafferent joint acceleration component [10]. From the DCIP EoM, a sensory model can be derived. Here, I repeat this EoM:

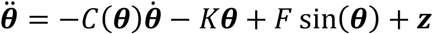

And I rewrite it as a function of acceleration instead of torque:

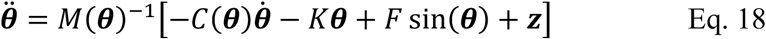

The right side of Eq. 18 can be linearized by computing the Jacobian matrices with respect to ***θ***, 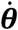 and ***z***, and evaluating them at the unstable fixed point **0**. I will denote these matrices by *K*_*stiff*_ , *C*_*damp*_ and *M*^−1^, and they correspond to, respectively, stiffness, damping and mass moment of inertia. Importantly, *K*_*stiff*_ equals *M*^−1^(*F* − *K*) (see Eq. 18) and thus specifies the combined contribution of gravity and joint-intrinsic stiffness. The linearized EoM can then be written as follows:

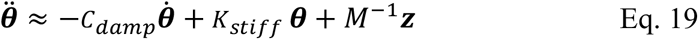

I start from Eq. 19 to derive an expression for the exafferent acceleration feedback provided by muscle spindles under fusimotor control [10]:

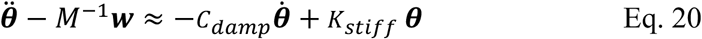

In Eq. 20, I have replaced ***z*** (noisy control signal sent to the mechanical system) by ***w*** (noisy fusimotor input to the muscle spindles) to allow for a different noise component in the two signals. This is in line with the simulations, in which there was a 50% shared variance in ***z*** and ***w***. The term *M*^−1^***w*** approximates the reafferent joint acceleration and its subtraction from the total acceleration 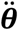 cancels this term from the spindle output. Eq. 20 can be written in matrix notation as follows:

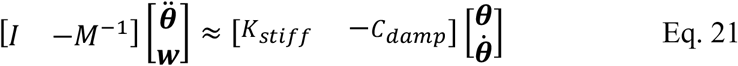

The coefficient matrices on the left and the right side of Eq. 21 are, respectively, the green and the blue matrix *C*_*Sens*_ and *C*_*Comp*_ in Fig. 2B. Thus,

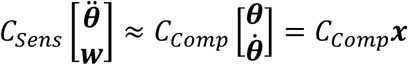

This shows that there exists a linear combination of the sensory feedback variables 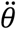 and ***w*** that equals a linear combination of the state variables ***x***. The matrix product *C*_*Comp*_***x*** is a sensory internal model.

### Simulations of standing balance control

#### The simulation algorithm

I now describe how to simulate the model without SMD in Fig. 2B. I will also mention the few adaptations that are required for a nonzero SMD, but I will not repeat the description of the SMD compensation mechanisms in the main text. I first describe the simulation of the mechanical system, the nonlinear DCIP. For this, I used the Matlab function ode45, which is based on an explicit Runge-Kutta (4,5) formula [86]. This function solves the DCIP EoM in state space form, which follows from Eq. 19:

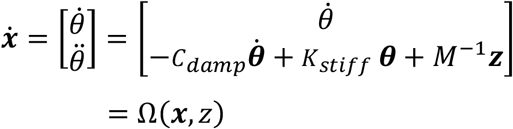

Simulating a mechanical system in discrete time requires a solution of its EoM. In fact, ***x***_*k*+1_ is obtained as the outcome of the function ode45 with input ***x***_*k*_ and ***z***_*k*_. And 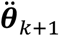 is obtained from Eq. 18 with input ***x***_*k*+1_ and ***z***_*k*_. For simulating a mechanical system with a SMD, ***z***_*k*_ is replaced by ***z***_*k*−*h*_, in which *h* is the SMD as a multiple of the discretization time step.

Exafferent acceleration feedback ***y***_*k*+1_ is computed by combining the acceleration output of the mechanical system 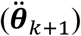 with the output of the fusimotor system (***w***_*k*+1_), and the pure sensory noise (***s***_*k*+1_):

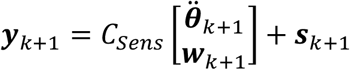

The computational system uses the following linear approximation of Ω(***x***, *z*):

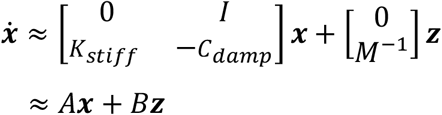

I assume that the mechanical system is deterministic, and therefore the motor noise *m* enters the mechanical system via the control signal ***z***: *Bz* = *B*(*u* + *m*) = *Bu* + *Bm*. Thus, the noise that is added to the mechanical system (*Bm*) is a linear function of the noise that is added to the control signal.

The computational system implements optimal control for the following linear dynamical system:

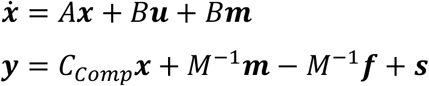

with Gaussian motor (***m***), fusimotor (***f***), and sensor (***s***) noise. The forward model in this system has a non-delayed input ***u***. To generalize this forward model to delayed input, I must express it explicitly as a function of time *t*: the forward model with non-delayed input involves ***u***(*t*), and the one with delayed input involves ***u***(*t* − *SMD*). This has only minor implications for the discrete time simulation algorithm: this algorithm must keep a memory buffer of efference copies ***u***(*t*) and select the appropriate one.

The optimal control problem is a LQG and its solution is given by a linear differential equation that governs the state estimates 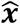:

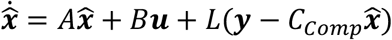

in which the control signal ***u*** equals 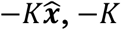 is the LQR gain and *L* is the Kalman gain [9]. Because of the definition of the motor and the fusimotor noise, there are a few minor adaptations to the usual way of calculating the Kalman gain: (1) the system noise covariance scaling matrix equals *B*var(***m***)*B*^*t*^ and thus depends on *B*, (2) the sensor noise covariance scaling matrix contains a term that depends on the fusimotor noise (i.e., 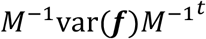), and (3) because the motor (***m***) and the fusimotor (***f***) noise can be correlated, this also holds for the system and the sensor noise.

#### Noise amplitude settings

To simulate DCIP dynamics under closed-loop feedback control, one must add noise. I set the noise amplitudes such that the effects of motor and pure sensor noise on the sensory feedback are equal. As a common scale for the effects of these three noise types, I use the noise variance of the sensory feedback ***y***_*k*_: if the noise source is only sensory, then *var*(***y***_*k*_) = *var*(***s***_*k*_), and if the noise source is only motor, then

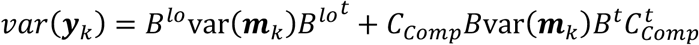

The matrix *B* is the discrete time version of the corresponding input matrix *B* in the continuous time linear dynamical system in Eq. 21. And the matrix *B*^*lo*^ is the lower half of the discrete time input matrix *B*. The details of the derivation are given elsewhere [10].

In the simulations, I start from a noise scaling variable *σ* and (1) set *var*(***s***_*k*_) = *σI*, in which *I* is an identity matrix of the appropriate dimension, (2) scale *var*(***m***_*k*_) such that the expression on the right side of Eq. 12 equals *σI*, and (3) set *var*(***f***_*k*_) = *var*(***m***_*k*_). I set the covariance *cov*(***m***_*k*_, ***f***_*k*_) such that the shared variance of ***m***_*k*_ and ***f***_*k*_ is 50%.

The variance of the noise terms *B****m***_*k*_ (for the state equation, but also fed into the output equation) and −*B*^*lo*^***f***_*k*_ + ***s***_*k*_ (for the output equation only) and their covariance are used as weights in the calculation of the Kalman gain *L*. This Kalman gain is a part of the computational system, and we thus implicitly assume that the CNS learns the sensor and the motor noise amplitudes from experience.

#### Physical and biophysical parameter settings

The simulations depend on several physical parameters that were assigned realistic and/or empirical values. I now list these parameters. First, the DCIP is specified by realistic and/or empirical values for its parameters:

1. Upper and lower body length: 0.75, resp., 1.1 m.
2. Upper and lower body mass: 50, resp., 35 kg.
3. Gravitational constant *g* = 9,8066.
4. Ankle stiffness = 493.47 Nm/rad. This number is based on measurements by [37], demonstrating that the ankle stiffness is 64% of the critical ankle stiffness (i.e., the ankle stiffness that would keep a standing person upright without active control).
5. Hip stiffness = 117.68 Nm/rad. This number is obtained in the same way as the ankle stiffness, assuming that the 64% also applies to the hip: hip stiffness is 64% of the critical hip stiffness.
6. Ankle damping = 30 Nm/(rad./s). This number was estimated by [37].
7. Hip damping = 19.10 Nm/(rad./s). This number was set such that the hip and the ankle damping ratios are equal. The hip damping was obtained by correcting the ankle damping for the different time constants of the two joints.

Second, for the simulations to be realistic, the computed ankle and hip torque must be less than the maximum voluntary contraction (MVC) of the muscle groups that must generate these two torque. Therefore, the output of the computational system is truncated at empirically determined MVC of these two muscle groups. For the ankle torque, I use plantarflexion MVC because humans prefer a leaned forward position [4]. In a study with 20 participants, the mean plantarflexion MVC was estimated to be 195 Nm [87]. To compute the hip torque MVC, I start from a setup involving a Roman chair in horizontal position on which a person lies sideways in a stretched position acting against gravity. The hip torque MVC is computed as the torque over the antero-posterior axis that is required to keep this challenging position. Using the length and mass parameter values in the previous paragraph, this MVC was estimated to be 184 Nm.

#### LQR gain weight matrices

The LQR gain −*K* is the result of an optimization that involves an objective function specified by two weight matrices: a 4-by-4 precision matrix *Q* and a 2-by-2 energetic cost matrix *R*. These weights are set such that the precision (*Q*-dependent) and the energetic cost (*R*-dependent) component have an equal contribution to the objective function. I use the maximum metric to compute matrices *Q* and *R* that weight precision and energetic cost equally: *Q* and *R* are scaled such that 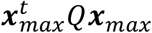 equals 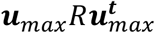.

The precision matrix *Q* has a block-diagonal structure diag(*Q*^*pos*^, *Q*^*vel*^) with *Q*^*pos*^ a 2-by-2 matrix that corresponds to the two angular position state variables (for upper and lower body) and *Q*^*vel*^ a 2-by-2 matrix that corresponds to the two angular velocity state variables. The matrices *Q*^*pos*^ and *Q*^*vel*^ are set such that the position and the velocity component have an equal contribution to the objective function: *Q*^*pos*^ and *Q*^*vel*^ are scaled such that 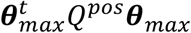 equals 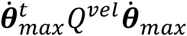. For the calculation of ***x***_*max*_, I started from the critical CoG angular position interval, which is the interval in which the CoG must stay such that the center of pressure does not leave the area of support. I calculated this interval from the ankle position, the foot length, and the CoG for a configuration with identical upper and lower body angular positions (no hip flexion of extention). I found this interval to be [-0.0758, 0.2293] rad., which spans a width of 0.3051 rad. Thus, ***θ***_*max*_ = [0.3051,0.3051]. Next, I calculated the maximum angular velocity from the maximum body sway frequency, which was estimated to be 1.875 Hz in [53]. From this maximum frequency, I calculated a maximum angular velocity of 0.3655 rad./sec. 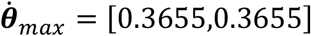. These values are sufficient to scale *Q*^*pos*^ and *Q*^*vel*^ such that the position and the velocity component have an equal contribution to the objective function.

The energetic cost matrix *R* is a diagonal 2-by-2 matrix, and it is scaled such that it has an equal contribution to the objective function as the precision (*Q*-dependent) component: 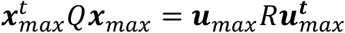. The maximum torques were inferred from the MVCs at the ankle and the hip joint, as described in an earlier paragraph: ***u***_*max*_ = [195,184] Nm.

#### Optimal gain scaling

In this paper, I applied optimal global scaling of the output gain: replace −*K* (the LQR gain for SMD=0) by −*αK*, in which *α* minimizes the LQR objective function. The minimization was performed using the Golden Section line search algorithm. As a part of this minimization routine, a function was called that computed a Monte Carlo estimate of the LQR objective function. This Monte Carlo estimate was computing the average LQR objective function over 10 simulation runs of 60 sec. each. Crucially, these simulations were ran with a nonzero SMD, and the outcome of the minimization is therefore specific for that SMD.

### The benchmark double pendulum bicycle model

The benchmark double pendulum (BDP) is based on three ideas. The first idea is to follow the approach of [55] and derive linearized EoM for a bicycle with the rider’s lower body rigidly attached to the rear frame and no upper body. These linearized EoM depend on a number of constants, and I chose these constants such that (1) the front frame is as similar as possible to the self-stable benchmark bicycle model described by [55]. The second idea is to model the interactions between the upper body and the rear frame (which includes the lower body) by the linearized EoM of the double compound pendulum, similar to [88]. Finally, the third idea is to first derive the BDP EoM without stiffness and damping terms, and to add these terms only in the last step.

The approach of [55] involves a method to calculate the defining matrices of linearized EoM of the following type:

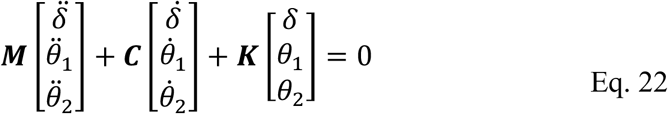

The matrices ***M, C*** and ***K*** are functions of several constants (angles, lengths, masses, mass moments of inertia, gravitational acceleration, speed) that characterize the bicycle components and the internal forces that act on them. It is straightforward to express the EoM in state space form by rearranging the terms in Eq. 22.

[55] only derived linearized EoM for bicycles with a rider that was rigidly attached to the rear frame. Thus, the upper body angular position *θ*_2_ is absent from their EoM. This missing component can be added by linearizing the DCIP EoM which models the interactions between *θ*_1_ and *θ*_2_. Schematically, each of the matrices ***M, C*** and ***K*** is composed as follows:

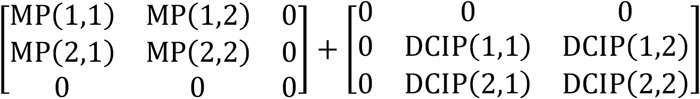

in which MP denotes matrix elements taken from Meijaard, Papadopoulos et al [55], and DCIP denotes matrix elements of DCIP EoM. The MP calculations were performed by means of the Matlab toolbox Jbike6 [89], in which I entered the constants for a bicycle with the rider’s lower body rigidly attached to the rear frame and no upper body. This produced the constants MP(*i*, *j*) (*i*, *j* = 1,2) for ***M, C*** and ***K***.

I now model the interactions between the upper and the lower body by the linearized DCIP EoM. I model the rear frame as a part of the lower body. I start from the following nonlinear EoM:

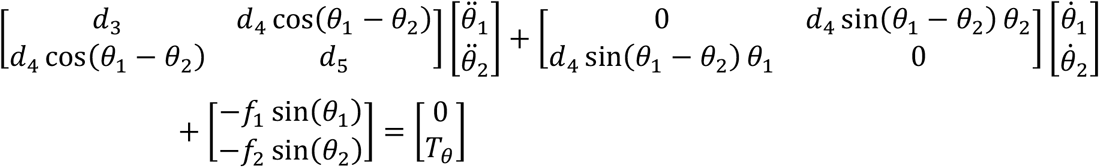

These EoM include the hip torque input *T*_*θ*_, but the ankle torque is set equal to 0. The constants *d*_3_, *d*_4_ and *d*_5_ contain elements that must be added to the matrix *M*, and the constants *f*_1_ and *f*_2_ contain elements that must be added to the matrix ***K***. These constants are defined as follows:

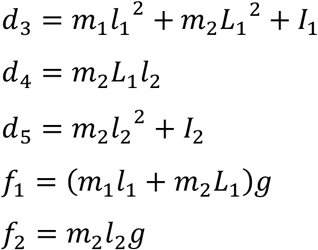

The constants *m*_1_, *L*_1_, *l*_1_ and *I*_1_ are, respectively, the mass, the length, the CoG (*L*_1_/2) and the mass moment of inertia of the DCIP’s first rod, which represents the rear frame and the rider’s lower body. The constants *m*_2_, *L*_2_, *l*_2_ and *I*_2_ are defined in the same way, but now for the second rod, which represents the rider’s upper body. Finally, *g* is the gravitational constant.

I evaluate these EoM at *θ*_1_ = *θ*_2_ and replace sin(*x*) by its linear approximation near 0: sin(*x*) ≈ *x*. This results in

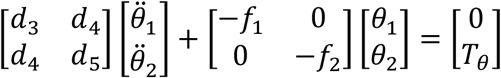

I will use the notation DCIP(*i*, *j*) (*i*, *j* = 1,2) to denote these elements. For *M*, the following elements are added:

- DCIP(1,1) = *m*_2_*L*_1_^2^
- DCIP(1,2) = DP(2,1) = *d*_4_ = *m*_2_*L*_1_*l*_2_
- DCIP(2,2) = *d*_5_ = *m*_2_*l*_2_^2^ + *I*_2_

And for ***K***, the following elements are added:

- DCIP(1,1) = *m*_2_*L*_1_*g*
- DCIP(2,2) = −*f*_2_ = −*m*_2_*l*_2_*g*

Finally, I added stiffness and damping terms to, respectively, ***K*** and *C*. As described in the next paragraph, these terms affect the dynamics of the steering and the hip joint.

Compared to the ankle joint in the DCIP model for standing balance control, much less is known about the stiffness and damping of the steering and hip joint in the BDP. These are not joints in the strict biomechanical sense because they involve more than the interface between two bones; the steering joint involves both the arms and part of the upper body, and the hip joint involves both the hip joint (head of femur and acetabulum) and the lumbosacral joint (lumbar spine and sacrum). For the steering joint, I calculated the stiffness from an empirically determined time constant, as described in [57]. And for the hip joint, I chose a stiffness coefficient such that the elastic force was 10 percent of the average (over upper and lower body) gravitational force; this allowed the upper and the lower body to fall with different accelerations. Evaluating the control performance using different damping ratios for the two joints (steering and hip) demonstrated robustness with respect to this parameter; control was possible over a range of four orders of magnitude around the critical damping ratio of 1 (i.e., for damping ratios between 0.01 and 100). I set the damping ratio for both joints at 20, which is a strongly overdamped system.

### Simulations of bicycle balance control

Except for the definitions and values of the physical and biophysical parameters, the simulation algorithm is identical to the one for standing balance control. The dimensionality of the state space representation increases from 4 to 6, but this does not affect the operations that must be performed to simulate the trajectories of the state variables.

Compared to standing balance control, torque limits are much less important in bicycle balance control. This is because (1) turning the handlebars requires very little torque, and (2) upper body position can also be controlled by the centrifugal torque, and thus not only depend on muscular torque. This was clear from the simulations, which produced much smaller torque values for bicycle than for standing balance control.

## Footnotes

1 The stability of this discrete time approximation in the context of closed loop control has been a topic of debate, and it has been shown that a low-pass filter of the predictions 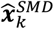 effectively deals with stability problems [14,17]. This debate was in the context of full-state feedback and therefore did not involve a state observer. In my simulations, I did not encounter stability problems and therefore do not consider low-pass filtering.

